# Mechanosensitive hormone signaling promotes mammary progenitor expansion and breast cancer progression

**DOI:** 10.1101/2022.04.19.487741

**Authors:** Jason J. Northey, Yoshihiro Yui, Mary-Kate Hayward, Connor Stashko, FuiBoon Kai, Janna K. Mouw, Dhruv Thakar, Jonathon N. Lakins, Alastair J. Ironside, Susan Samson, Rita A. Mukhtar, E. Shelley Hwang, Valerie M Weaver

**Author notes:** **Corresponding Author:** Valerie M. Weaver, Center for Bioengineering and Tissue Regeneration, Department of Surgery, University of California, San Francisco.

## Abstract

Tissue stem-progenitor cell frequency has been implicated in tumor risk and progression. Tissue-specific factors linking stem-progenitor cell frequency to cancer risk and progression remain ill defined. Using a genetically engineered mouse model that promotes integrin mechanosignaling with syngeneic manipulations, spheroid models, and patient-derived xenografts we determined that a stiff extracellular matrix and high integrin mechanosignaling increase stem-progenitor cell frequency to enhance breast tumor risk and progression. Studies revealed that high integrin-mechanosignaling expands breast epithelial stem-progenitor cell number by potentiating progesterone receptor-dependent RANK signaling. Consistently, we observed that the stiff breast tissue from women with high mammographic density, who exhibit an increased lifetime risk for breast cancer, also have elevated RANK signaling and a high frequency of stem-progenitor epithelial cells. The findings link tissue fibrosis and integrin mechanosignaling to stem-progenitor cell frequency and causally implicate hormone signaling in this phenotype. Accordingly, inhibiting RANK signaling could temper the tumor promoting impact of fibrosis on breast cancer and reduce the elevated breast cancer risk exhibited by women with high mammographic density.

**Summary:** Elevated mechano-signaling and matrix stiffness promote progesterone and RANK mediated expansion of mammary progenitors and breast cancer risk and progression.

## INTRODUCTION

The frequency of stem-progenitor cells and the level of cell proliferation within a tissue correlate with overall level of risk to malignancy^1–3^. The frequency of cancer cells with a stem-progenitor phenotype also dictates tumor aggression and may foster metastasis and confer treatment resistance^4–8^. Populations of stem-progenitor cells within a tumor may reflect the expansion of genetically modified resident stem cells or the transdifferentiation of resident tumor cells towards a stem-progenitor-like phenotype^9,10^. Nevertheless, what tissue-specific factors regulate stem-progenitor cell frequency in normal and malignant tissues remains poorly understood.

The extracellular matrix (ECM) influences tissue development and homeostasis^9,11^. ECM properties also regulate the stem cell niche and its stiffness modulates stem cell growth, survival and tissue-specific differentiation^12–15^. Consistently, the ECM is altered in tumors and the most aggressive tumors, which are usually metastatic and harbor the highest frequency of stem-progenitor cells, are often the most fibrotic and have the stiffest stroma with the greatest amount of cross-linked fibrillar collagen^16–19^. Indeed, a stiff ECM promotes the epithelial-to-mesenchymal transition (EMT) of epithelial cells, and tumor cells that have undergone an EMT are enriched for the expression of stem-progenitor markers and exhibit stem-progenitor-like phenotypes^20–22^. Although it remains unclear how, these findings do suggest that the stiffness of the tissue ECM could modulate the number of stem-progenitor cells.

A stiff ECM modifies cell and tissue behavior, in part, by regulating the context of cellular signaling to influence gene expression^9,11,23^. Specifically, a stiff ECM fosters the assembly of integrin focal adhesions which potentiate growth factor dependent ERK and phosphatidylinositol 3-kinase (PI3K) signaling^24–28^. ECM stiffness also amplifies cytokine-stimulated G-Protein-Coupled Receptor (GPCR) activity^14,19,29–31^, permits morphogen-induced Notch and Wnt activation^14,15,26^ and facilitates β-catenin signaling^32^. Thus, we speculated that a stiff ECM stroma could expand tissue-resident stem-progenitor frequency and fate by altering the context of cell signaling to modify gene expression. To test this theory, we opted to study the mammary gland. The mammary gland is a unique tissue that undergoes development in the adult organism where the growth, survival, invasion and differentiation of the mammary epithelium and stem-progenitor cells are tightly regulated by a circuit of hormones including estrogens, progesterone and prolactin. Progesterone in particular has been implicated in regulating the expansion of the stem-progenitor epithelial pool in the breast^33–35^. Progesterone supplementation has also been linked to increased breast cancer risk in postmenopausal women^36–40^, and has been causally linked to breast tumor progression and aggression in experimental models^41–44^. Indeed, estrogen, both stimulates progesterone expression and regulates mammographic density (MD), with high MD conferring an elevated lifetime risk of breast cancer ^45–47^. Moreover, high MD breast stroma is significantly stiffer than the low MD breast stroma^48,49^.

This led us to hypothesize that a stiff ECM regulates progesterone activity to expand the pool of mammary epithelial progenitors in both normal tissue and in fibrotic stiff breast tumors. We used transgenic, orthotopic and patient derived models of breast cancer in combination with organoid and spheroid models to test this theory. Our studies revealed that a stiff ECM and elevated integrin mechanosignaling potentiate progesterone-induced RANK signaling to expand the pool of mammary epithelial stem-progenitors in both normal tissue and in highly fibrotic breast tumors. The findings suggest targeting RANK signaling might reduce breast cancer risk in women with high MD and have potential therapeutic implications in patients with highly fibrotic breast cancers.

## RESULTS

### Integrin mechanosignaling promotes metastasis by inducing a mesenchymal phenotype

Tissue fibrosis is a feature of breast cancers that associates with tumor aggression. Fibrotic tumors have increased levels of cross-linked ECM that stiffen the stroma and elevate tumor cell integrin mechanosignaling^16–18,50^. A stiff ECM promotes the expression of mesenchymal-associated genes and drives the mesenchymal-like invasive phenotype of tumor cells in culture and the metastatic behavior of tumors *in vivo* ^25,26,51–53^. In a cohort of 167 breast cancer patients with basal-like tumors, those with upregulation of integrin/EMT related genes were significantly more likely to fail to undergo a complete pathologic response (pCR) after neoadjuvant chemotherapy as compared to those without upregulation of these genes (Fig. 1a)^54^. To investigate whether a causal role exists between ECM stiffness, integrin-mechanosignaling and tumor aggression we engineered mice to conditionally express a mutant human β1 integrin (V737N) that promotes focal adhesion assembly and signaling independent of substrate stiffness. V737N-β1 integrin transgene expression was targeted to mammary epithelial cells (MECs) using MMTV-Cre (MMTV-Cre/β1-V737N^fl/-^; “V737N”) (Extended Data Fig. 1a)^26^. We confirmed V737N-β1 expression within the mammary epithelium and noted its preferential enrichment in the basal compartment using flow activated cell sorting (FACS) and immunofluorescence staining to monitor EGFP levels that were coupled to the transgene using an IRES element (Extended Data Fig. 1b,c). We then crossed the MMTV-Cre/β1-V737N^fl/-^ mice with MMTV-Neu mice to generate Neu-induced tumors with (MMTV-Neu^+/-^/MMTV-Cre^+/-^/β1-V737N^fl/-^ or “V737N^Neu^”) and without (MMTV-Neu^+/-^/MMTV-Cre^-/-^/β1-V737N^fl/-^ or “Neu”) V737N-β1 (Fig. 1b). After demonstrating that the V737N^Neu^ mice maintained robust expression of the mutant β1 integrin in their mammary epithelium (confirmed using a human specific antibody to β1 integrin; Extended Data Fig. 1d) we monitored the mice for tumor latency, phenotype and metastasis as compared to age-matched Neu mice. Immunofluorescence staining revealed that phospho-ERK, phospho-AKT substrates, phospho-FAK (Y397) and active β1 integrin were increased in the V737N^Neu^ tumors, demonstrating that V737N-β1 expression successfully elevated integrin mechanosignaling (Extended Data Fig. 1e-h)^26^. Tumor latency was not appreciably different between the Neu and V737N^Neu^ mice (Extended Data Fig. 1i). We also failed to observe any changes in the rate of tumor outgrowth which was confirmed by our inability to detect any alterations in phospho-Histone-H3 staining by immunofluorescence between Neu and V737N^Neu^ mice (Extended Data Fig. 1j,k). Furthermore, tumor cell viability was unchanged as indicated by similar levels of cleaved-caspase 3 in the two experimental groups (Extended Data Fig. 1l). Nevertheless, and importantly, lung metastasis was profoundly increased in the V737N^Neu^ mice (60% V737N^Neu^ vs. 10% Neu; Extended Data Fig. 1m) with a 7-fold increase in the average number of lung lesions per animal (Fig. 1c). Lung lesions in V737N^Neu^ mice were also significantly larger than those in Neu mice (Fig. 1d). These findings suggest there were likely qualitative differences in the nature of the tumors that developed in the V737N^Neu^ mice as compared to the Neu mice.

**Fig. 1:**
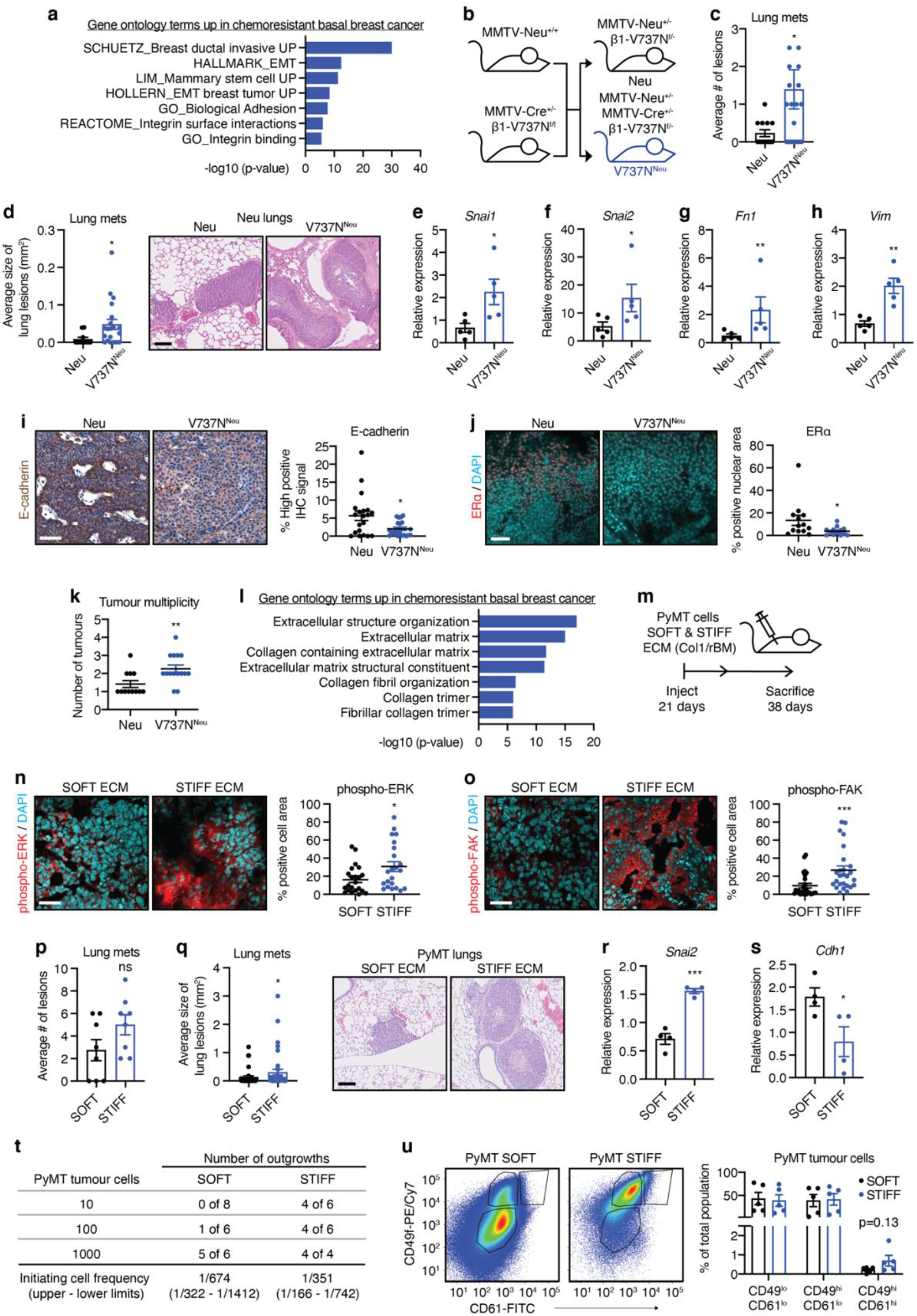
Mechanosignaling and extracellular matrix stiffness promote a mesenchymal tumor phenotype and mammary tumor initiation. **a**, GSEA of transcriptional profiles from the Nuvera study showing upregulated Epithelial-to-mesenchymal transition and integrin adhesion gene ontology terms associated with poor response to chemotherapy. **b**, Breeding schematic for Neu and V737N^Neu^ transgenic mice. **c**, Average number of lung metastases for Neu (n=13) and V737N^Neu^ (n=19) mice determined by histological analysis. **d**, (left) Average size of the metastatic lesions corresponding to the analysis in c. (n=13; Neu mice, n=19; V737N^Neu^ mice). (right) Representative H&E images of lung lesion size. Scale bar, 100 μm. **e-h**, qRT-PCR analysis of RNA isolated from Neu (n=5) and V737N^Neu^ (n=5) tumors showing the relative expression of the indicated mesenchymal genes. **i**, (left) Representative immunohistochemical staining of Neu (n=5) and V737N^Neu^ (n=5) paraffin tumor sections with an E-cadherin specific antibody. Scale bar, 50 μm. (right) Graph displaying quantification of average high-positive signal for each group. **j**, (left) Representative immunofluorescence staining of Neu (n=5) and V737N^Neu^ (n=5) frozen tumor sections with an ERα specific antibody. Scale bar, 25 μm. (right) Graph showing quantification of ERα-positive nuclear area. **k**, Dot plots of the average tumor multiplicity (number of tumors per animal) for Neu (n=13) and V737N^Neu^ (n=19) mice. **l**, GSEA of transcriptional profiles from the Nuvera study showing upregulated extracellular and collagen matrix gene ontology terms associated with poor response to chemotherapy. **m**, Schematic and timeline of syngeneic orthotopic implantation of PyMT tumor cells within SOFT (non-crosslinked, no L-ribose) and STIFF (crosslinked, +L-ribose) Col1/rBM hydrogels. **n**, (left) Representative immunofluorescence staining of SOFT (n=5) and STIFF (n=5) PyMT frozen tumor sections with an antibody specific to phospho-ERK. Nuclei are counterstained with DAPI. Scale bar, 50 μm. (Right) Graph showing quantification of average positive cell area. **o**, (left) Representative immunofluorescence staining using an antibody specific to phospho-FAK for samples from n. Scale bar, 50 μm. (right) Graph showing quantification of average positive cell area. **p**, Average number of lung metastases for SOFT (n=8) and STIFF (n=8) PyMT tumor bearing mice determined by histological analysis. **q**, (left) Average size of the metastatic lesions corresponding to the analysis in p. (right) Representative images of H&E-stained lung sections. Scale bar, 100 μm. **r-s**, qRT-PCR analysis of gene expression for the indicated genes using RNA isolated from SOFT (n=4) and STIFF (n=4) PyMT tumors. **t**, Tumor initiating cell frequencies were measured for SOFT and STIFF PyMT tumors (from a pool of three individual tumors each) by primary and secondary limiting dilution transplantation tumorigenesis assays in syngeneic mice. **u**, FACs analysis of tumors using CD24, CD49f and CD61 cell surface antigens to determine the percentage of tumor initiating cells from SOFT (n=5) and STIFF (n=5) PyMT tumors defined as CD49f^hi^; CD61^hi^. (left) Representative FACS dot plots for SOFT and STIFF PyMT tumors. (right) Graph displays average percentages calculated for the indicated cell populations from tumors in each group. All graphs are presented as mean +/- S.E.M. Statistical tests used were Mann-Whitney test (c, d, g i-k, n-q) and unpaired *t*-test (e, f, h, r, s, u), *P<0.03, **P<0.002, ***P<0.0002, ns=non-significant.

We next examined the histological phenotype of the tumors formed in Neu and V737N^Neu^ mice. A higher proportion of tumors in the V737N^Neu^ mice developed dense, highly packed tumors lacking luminal space, and displayed striking nuclear morphologies with increased nuclear pleomorphism and conspicuous mitotic activity, both components of pathologic grade, compared to the control Neu mice (5 of 8 versus 2 of 7 respectively; Extended Data Table 1). Consistently, V737N^Neu^ tumors expressed significantly higher transcript levels for several mesenchymal genes including *Snai1*, *Snai2*, *Fn1*, and *Vim* as compared to Neu controls (Fig. 1e-h). Immunohistochemistry (IHC) and immunofluorescence for the luminal epithelial markers E-cadherin and ERα demonstrated decreased protein expression (Fig. 1i,j) that aligned with quantitative reverse transcriptase-PCR (qRT-PCR) analysis of their corresponding mRNAs (Extended Data Fig. 1n,o). Perhaps most intriguingly, there was an almost two-fold increase in the number of tumors per animal (tumor multiplicity) in the V737N^Neu^ mice, suggesting elevated mechanosignaling might enhance the potential for tumor initiation (Fig. 1k). The data indicate that increased integrin-mechanosignaling could cultivate a mesenchymal-like cell phenotype that enhances the ability of mammary tumor cells to metastasize and that might additionally foster tumor initiation.

A stiff, fibrotic ECM increases integrin activity and focal adhesion assembly to enhance mechanosignaling. Indeed, ECM-related gene ontology terms were among the top hits upregulated in the chemoresistant patient tumors from the Nuvera study (Fig. 1l)^54^. To test whether a fibrotic ECM could promote breast tumor aggression and initiation by enhancing integrin signaling, we embedded MMTV-PyMT mouse derived tumor cells into the fat pads of syngeneic mice within collagen-I/reconstituted basement membrane hydrogels (Col1/rBM) with and without L-ribose meditated crosslinking to generate a “STIFF” (∼1700 Pascals) and a “SOFT” (∼600 Pascals) ECM stroma, respectively (Fig. 1m)^25,55^. Immunofluorescence staining for phospho-ERK, phospho-FAK and active β1 integrin demonstrated increased mechanosignaling in the tumor cells implanted within the STIFF ECM stroma, likely accounting for their modestly increased rate of growth (Fig. 1n,o and Extended Data Fig. 2a,b). In agreement with results from the V737N^Neu^ mice, the PyMT tumors in the STIFF ECM stroma gave rise to a higher frequency and larger lung metastasis (Fig. 1p,q and Extended Data Fig. 2c). qRT-PCR analysis confirmed that the PyMT tumors that developed within the STIFF stroma expressed higher levels of mesenchymal genes including *Snai2, Zeb1* and *Krt5* and lower levels of epithelial markers such as *Cdh1* (Fig. 1r,s and Extended Data Fig. 2d,e). Importantly, the PyMT tumor cells harvested from the STIFF ECM stroma demonstrated a greater tumor initiating potential as assessed by limiting dilution tumorigenesis assays when compared to the potential of tumor cells isolated from the SOFT ECM stroma (Fig. 1t and Extended Data Fig. 2f). FACS analysis of tumor cells isolated from the SOFT and STIFF ECM stroma revealed there was an increased frequency of a CD24^+^CD49^hi^CD61^hi^ pool of tumor initiating cells in the tumors that developed within the STIFF ECM stroma (Fig. 1u). The data extend and reinforce prior observations that a stiff, fibrotic ECM and elevated β1 integrin mediate mechanotransduction that induces a mesenchymal phenotypic shift in tumor cells that potentiates tumor initiation and aggression and that drives metastases.

### Tissue tension enhances tumor initiation and metastasis by increasing the number of human patient-derived tumor cells with a mesenchymal, progenitor-like phenotype

Human breast cancer progression and aggression associate with the presence of oriented, thick and stiffened extracellular collagens, particularly surrounding developing lesions and at the invasive front of the tumor^16,17,56^. Consistently, atomic force microscopy (AFM) confirmed a significant correlation between the stiff ECM stroma at the invasive front of HER2-positive human breast tumor tissue and high intensity immunofluorescence staining for active β1 integrin (Fig. 2a). We therefore examined whether a causal relationship exists between a stiffened stroma and human breast cancer aggression, and if this was mediated through elevated integrin-mechanosignaling. We implanted three independent HER2-positive breast cancer patient derived xenografts (PDXs) embedded within either a SOFT or a STIFF Col1/rBM ECM stroma into the mammary fat pads of host mice and monitored for impact on tumor phenotype and behavior (Fig. 2b)^57,58^. Immunofluorescence for phospho-ERK, phospho-FAK and active β1 integrin verified that a STIFF ECM stroma enhanced mechanosignaling in the cells of these human tumors relative to that observed in the tumor cells embedded within the SOFT ECM stroma (Fig. 2c,d and Extended Data Fig. 3a). We observed a modest but consistent increase in tumor outgrowth in all three independent HER2-positive PDX tumors (Extended Data Fig. 3b-d). The STIFF ECM stroma also increased the frequency and size of lung metastasis in two of the three PDXs; BCM-3963 and BCM-3143B (Fig. 2e-h and Extended Data Fig. 3e,f). Analysis of RNA-seq gene expression data (MSig database)^59,60^ generated from the three independent HER2-positive PDX models revealed a significant upregulation of genes in categories related to cell adhesion, EMT and stem/progenitor cells in the tumors that developed in the STIFF ECM stroma (Fig. 2i). Plotting fold change versus p-values (-log10) of the RNA-seq data illustrates several of the upregulated mesenchymal markers from the Hallmark_Epithelial_Mesenchymal_Transition category (Fig. 2j). PCR arrays validated the RNA-seq expression analysis and identified several additional EMT markers whose expression levels were increased towards that defined as a mesenchymal phenotype in the HER2-positive PDX tumors that developed within the STIFF ECM stroma (Fig. 2k-n and Extended Data Fig. 3g-l). All three independent HER2-positive PDX tumors also demonstrated higher tumor initiation when they were implanted within the STIFF ECM stroma (Fig. 2o-q). Furthermore, FACS analysis revealed that the tumors that developed within the STIFF ECM stroma harbored a significantly higher proportion of CD24^lo^CD44^hi^ or EpCAM^hi^CD49f^hi^ tumor initiating cells (Fig. 2r,s). These findings provide further evidence that a stiff ECM stroma stimulates integrin-mechanosignaling that in turn upregulates transcriptional programs that generate more mesenchymal-like tumor cells with enhanced capacity to initiate tumors, disseminate and form metastatic lesions.

**Fig. 2:**
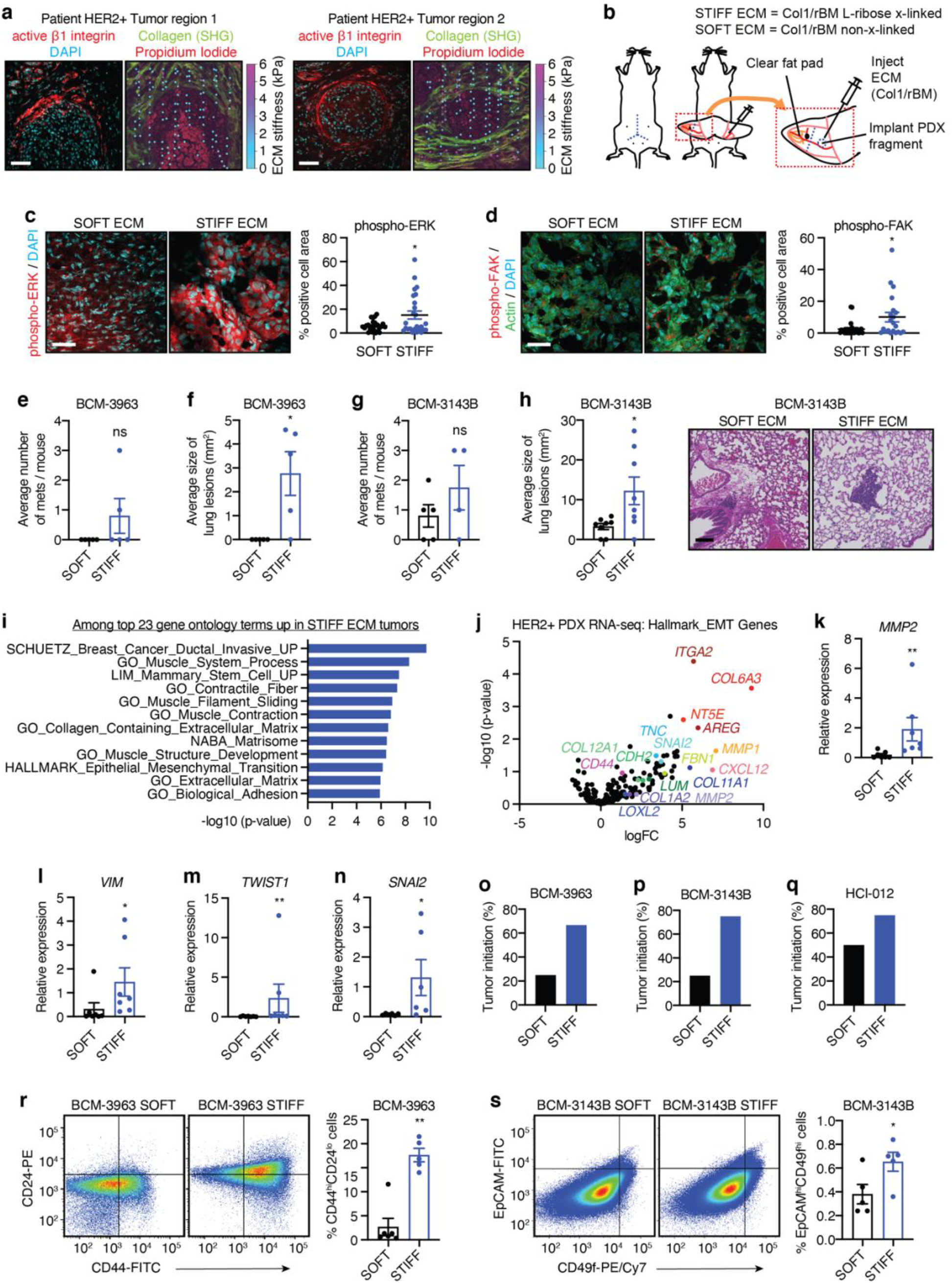
Extracellular matrix stiffness promotes a mesenchymal phenotype and aggression in human breast cancer. **a-b**, Sequential frozen sections from two regions of a HER2-positive breast cancer patient specimen were subjected to Atomic Force Microscopy (AFM) analysis, Second Harmonic Generation (SHG) imaging (collagen matrix), and immunofluorescence staining. (right) Sections analyzed by AFM were counterstained with propidium iodide to visualize nuclei and orient measurements. The same sections were used for SHG imaging by 2-photon microscopy (green) and AFM measurements (single indentations in a grid pattern) are overlayed as color-scaled dots (cyan-magenta). Scale bar, 80 μm. (left) Sequential sections were stained by immunofluorescence with an antibody to β1-integrin (active conformation; red) and nuclei were counterstained with DAPI (cyan). **b**, Schematic showing the strategy for implantation of HER2-positive patient-derived xenograft (PDX) breast cancer tissues in SOFT (Col1/rBM, no L-ribose) and STIFF (Col1/rBM, crosslinked with L-ribose) ECM stroma. **c**, (left) Representative images of immunofluorescence staining of frozen tumor sections from HER2-positive PDX tumors generated within a SOFT (n=6) and STIFF (n=6) ECM stroma with an antibody specific to phospho-ERK. Scale bar, 50 μm (right) Graph showing quantification of average phospho-ERK positive cell area for all HER2-positive PDX models combined. **d**, (left) Representative images of immunofluorescence staining as in c (SOFT; n=6, STIFF; n=6) using an antibody specific to phospho-FAK. Scale bar, 50 μm. (right) Graph showing quantification of average phospho-FAK positive cell area for all HER2-positive PDX models combined **e**, Average number of lung metastases for mice bearing BCM-3963 PDX tumors in SOFT (n=10) and STIFF (n=10) ECM stroma as determined by histological analysis. **f**, Average size of the metastatic lesions corresponding to the analysis in e. **g**, Analysis as in e for mice bearing BCM-3143B PDX tumors (SOFT; n=10, STIFF; n=10). **h**, Analysis as in f for mice bearing BCM-3143B PDX tumors (SOFT; n=10, STIFF; n=10). (right) Representative images of lung metastases for mice bearing BCM-3143B PDX tumors in SOFT and STIFF ECM stroma. Scale bar, 100 μm. **i**, Gene ontology terms from among the top 23 most significantly upregulated, using RNAseq data derived from all HER2-positive PDX tumors generated in SOFT (n=9) and STIFF (n=9) ECM stroma as above (n=3 for each PDX and condition). **j**, Scatter plot of p-value (-log10) vs. log fold change (logFC) for gene expression from the HALLMARK_epithelial-to-mesenchymal transition gene set for RNAseq data of HER2-positive PDX tumors developed in SOFT and STIFF ECM stroma **k-n**, qRT-PCR arrays designed to examine Epithelial-to-mesenchymal transition related gene expression were used to analyze RNA isolated from PDX tumors developed in SOFT (n=7) and STIFF (n=7) ECM stroma. Bar plots for the average relative expression of the indicated mesenchymal genes are displayed. **o-q**, Tumor initiation represented as the percentage of tumors established in mice with SOFT and STIFF ECM stroma from cryopreserved tumor fragments for each indicated HER2-positive PDX model. **r**, (left) Representative FACS plots for analysis of tumor initiating cell frequency in the PDX model BCM-3963 for tumors developed within SOFT (n=5) and STIFF (n=5) ECM stroma. (right) Graph showing the average percentage of CD44^hi^CD24^lo^ tumor initiating cells. **s**, Representative FACS plots for analysis of the PDX model BCM-3143B as in q (SOFT; n=5, STIFF; n=5). (right) Graph showing the average frequency of tumor initiating cells expressed as the percentage of EpCAM^hi^CD49f^hi^ cells. All graphs are presented as mean +/- S.E.M. Statistical tests used were Mann-Whitney test (d, f, k-m, r) and unpaired *t*-test (c, e, g, h, s), *P<0.03, **P<0.002, ***P<0.0002, ns=non-significant.

### High mechanosignaling promotes stem/progenitor-like activity in mammary epithelial cells

We next explored whether elevating integrin-mechanosignaling, per se, could expand stem-progenitor frequency in the normal mammary gland, and if so, how? We studied mammary gland development in 6- and 10-week-old non-tumor bearing V737N mice and compared our findings to non-tumor bearing control mice (CTL) (Fig. 3a). Immunofluorescence staining for human β1 integrin, phospho-ERK, phospho-FAK and phospho-p130CAS demonstrated elevated levels in the MECs of the mice expressing the V737N β1 integrin, consistent with increased activity of integrin-mechanosignaling in the mammary epithelium (Extended Data Fig. 4a-c)^26^. Gross examination of mammary glands from 6- and 10-week-old V737N mice compared to age matched CTL mice further revealed that the mammary glands of V737N mice were larger and heavier (Extended Data Fig. 4d-f). Further histological examination of the mammary glands of 6-week-old mice revealed an increase in the number and size of terminal end buds in V737N mice (Fig. 3b). We also observed precocious primary, secondary and tertiary ductal branching in the V737N mice at both 6- and 10-weeks of age (Fig. 3c and Extended Data Fig. 4g,h). Concordant with these data, proliferation as assessed by immunofluorescence staining for phospho-Histone H3 was elevated in the V737N β1 integrin expressing MECs compared to CTL MECs (Extended Fig. 4i,j). Examination of H&E-stained cross-sections of the mouse mammary ducts, revealed that the basal/myoepithelial layer of MECs was thicker in the V737N β1 integrin expressing mammary glands (Fig. 3d). Immunofluorescence staining for lineage specific cytokeratin markers (Luminal, K8 and Basal, K14) at both 6- and 10-weeks of age (Fig. 3e) confirmed a striking thickening of the basal layer as revealed by an increase in cells staining positive for K14. Subsequent FACS analysis of isolated MECs clearly demonstrated a higher proportion of basal to luminal MECs in V737N mammary glands indicating that the basal compartment was significantly expanded (Fig. 3f). The data suggest that high integrin-mechanosignaling fosters the expansion of basal MECs in the mammary gland.

**Fig. 3:**
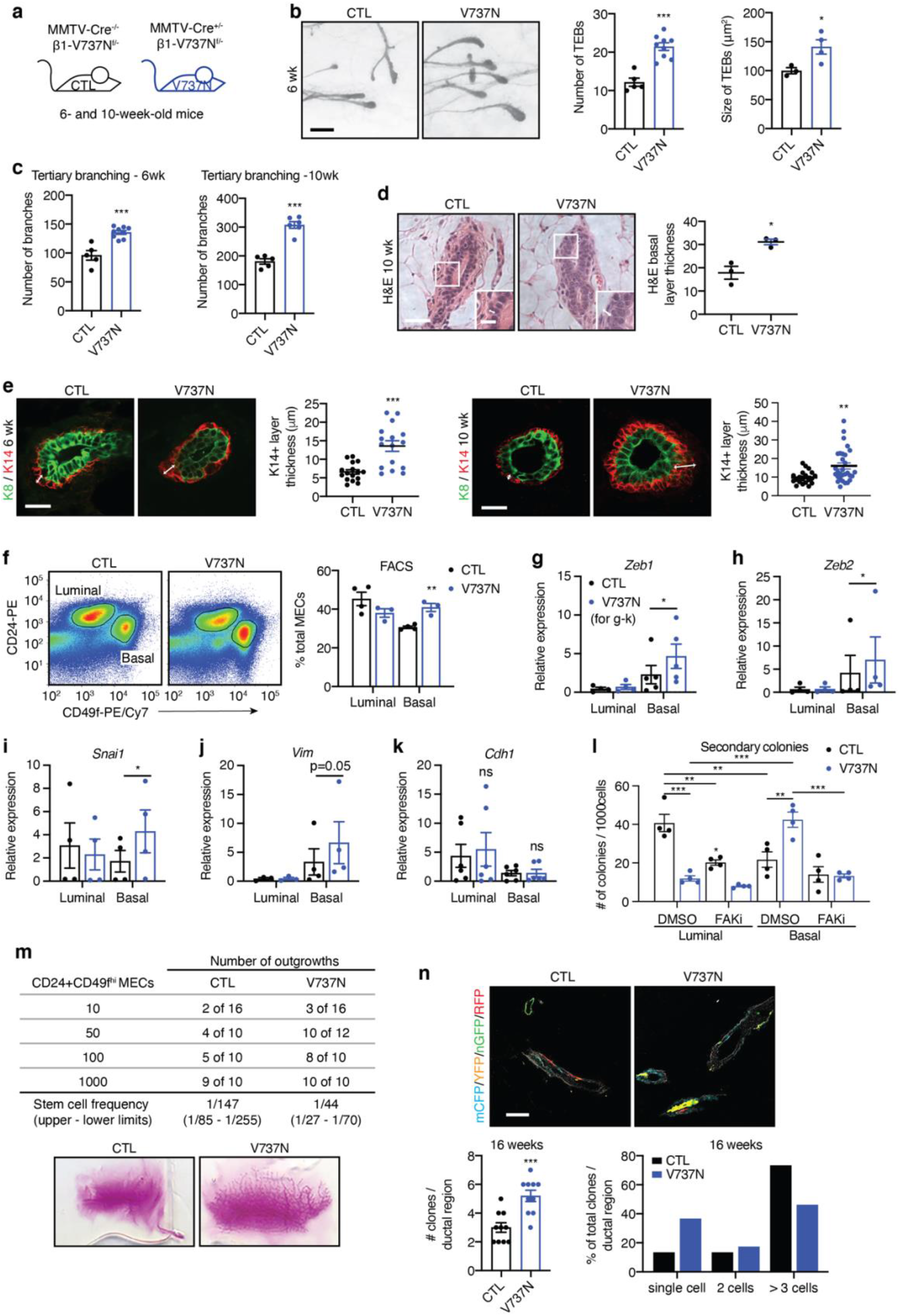
High mechanosignaling fosters stem/progenitor activity in mammary epithelial cells. **a**, Cartoon illustrating the genotypes that reflect wildtype (CTL) and V737N-β1 integrin (V737N) expression in MECs in mice. **b**, (left) Representative images of H&E-stained mammary gland wholemounts from CTL and V737N mice. Scale bar, 400 μm. (middle and right) Graphs showing the average number and size of ductal terminal end buds (TEBs) (CTL; n=3-5, V737N; n=3-8 mice). **c**, Graphs of the average number of tertiary ductal branching observed for CTL and V737N mice at 6 (left) and 10 (right) weeks of age (CTL and V737N; n=5-9 mice). **d**, (left) Representative H&E-stained histological sections of mammary glands from 10-week-old CTL and V737N mice. Scale bar, 50 μm (main image and inset). (right) Graph showing quantification of the average thickness of the basal/myoepithelial layer (CTL and V737N; n=3 mice). **e**, Representative immunofluorescence of luminal and basal keratins (K8; green and K14; red) for frozen sections of mammary glands from CTL and V737N mice (n=5 mice) at 6- (far left) and 10- (middle right) weeks of age. Scale bar, 50 μm. Graphs showing quantification of the average thickness of K14-postive layers for mice at 6- (middle left) and 10- (far right) weeks of age. **f**, (left) Representative FACS plots showing analysis for mammary epithelial lineages from CTL and V737N mice using CD24 and CD49f surface antigens. (right) The relative average percentages of luminal (CD24^+^; CD49f^lo^) and basal (CD24^+^; CD49f^hi^) populations are plotted (n=4 independent isolations each for CTL and V737N mice). **g-k**, RNA was isolated from luminal and basal lineage MECs sorted from CTL and V737N mice (n=4-6 independent isolations) as in f. Graphs showing relative expression for the indicated genes are displayed. **l**, MECs sorted from CTL and V737N mice as in f were subjected to rBM colony formation assays to assess luminal and basal progenitor activity (n=4 independent isolations). The bar graph shows secondary colony formation in the presence of vehicle (DMSO) or FAK-inhibitor (FAKi; PND1186) **m**, Limiting dilution transplantation assays were performed with basal (CD24^+^; CD49f^hi^) MECs sorted as in f from CTL and V737N mice. Stem cell frequency was calculated from the number of mammary gland repopulating events. (bottom) Representative images of carmine alum-stained mammary gland transplants of CTL and V737N MECs **n**, Lineage tracing of CTL and V737N mammary glands using tamoxifen (single dose of 1.5mg per mouse) induced V737N-β1 and Confetti reporter expression at 3-4 weeks of age in K5-positive epithelial cells (K5/creERT2) (n=3 for CTL and V737N mice). (top) Representative images of fluorescent protein-labelled cells 16 weeks post tamoxifen induction. Scale bar, 50 μm. Frequency of distinct clones per ductal region (bottom left) and the relative percentage of single cell, two-cell, and multicellular clones (bottom right) are represented as bar plots. All graphs are presented as mean +/- S.E.M. Statistical tests used were unpaired *t*-test (b-e), two-way ANOVA (with Tukey’s multiple comparisons test) (f, l) and paired two-way ANOVA (with two-stage linear step-up procedure of Benjamini, Krieger and Yekutieli for multiple comparisons test) (g-k). *P<0.05, **P<0.005, ***P<0.0005, ns=non-significant.

To determine whether the higher frequency of basal MECs reflected an increase in progenitor abundance or if high mechanosignaling induced the transdifferentiation of MECs, we utilized FACS to isolate luminal and basal MECs from CTL and V737N mice. Gene expression analysis for known luminal and basal markers (*Krt8*, *Krt18*, *Krt5*, *Krt14* and *Muc1*) confirmed efficient sorting of the two different MEC populations (Extended Data Fig. 5a-e). Further qRT-PCR analysis revealed that the mesenchymal markers *Zeb1*, *Zeb2*, *Snai1* and *Vim* were elevated in V737N-β1 expressing basal MECs as compared to CTL MECs, while levels of the luminal marker *Cdh1* remained largely unchanged (Fig. 3g-k). Given that putative mammary stem-progenitor cells have been proposed to reside within the basal MEC compartment, we next investigated whether mechanosignaling enhanced the functional stem-progenitor behavior of the isolated MECs. FACS-sorted luminal and basal MECs were subjected to colony formation assays in rBM hydrogels. Although primary mammosphere colony efficiency was identical between the isolated CTL and V737N luminal and basal MECs (Extended Data Fig. 6a), secondary colony formation revealed a significant increase in mammary colony formation in the V737N basal MEC population and a corresponding decrease in V737N luminal progenitors (Fig. 3l). Importantly, this effect was abolished when the mammary colonies were treated with a FAK inhibitor (FAKi; PND-1186) consistent with stem-progenitor population self-renewal being dependent upon the elevated β1 integrin mediated mechanosignaling (Fig. 3l). Limiting dilution transplantation assays (LDTAs) in which the ability of the basal (CD24^+^CD49f^hi^) MEC populations isolated from the CTL and V737N β1 integrin expressing MECs to reconstitute the mammary gland was assessed by injecting the cells into the cleared fat pads of syngeneic host mice, revealed a nearly 3-fold increase in stem-progenitor cell frequency for the V737N MECs over the CTL MECs (Fig. 3m). To more rigorously test whether increasing integrin-mechanosignaling enhanced stem-progenitor cell frequency we next used a developmental lineage tracing strategy to examine the clonal dynamics of MECs by monitoring clonal diversity during branching morphogenesis^61^. For this purpose, the basal K5 promoter was used to activate lineage tracing and V737N expression in mouse MECs (K5-Cre/ERT2^+/-^/Brainbow2.1^+/-^/V737N^fl/-^ mice) with one low dose of Tamoxifen at puberty (3-4 weeks old). A 2-week trace displayed little differences in clonal frequency or size (number of cells/clone) except for a slight increase observed in the frequency of multicellular clones in V737N-β1 expressing glands that may reflect a propensity for increased proliferation (Extended Data Fig. 6b,c). However, a significant increase in clonal frequency was observed at 16 weeks post tamoxifen induction in the mammary glands of V737N mice as compared to CTL mice (Fig. 3n). Moreover, CTL mammary glands showed a reduction in single cell clones concomitant with an increase in multicellular clones. By comparison, single cell clones persisted in mammary glands of the V737N mice suggesting their stem-progenitor potential is likely maintained over longer periods of time (Fig. 3n).

### RANKL is required for mechanosignaling induced mammary stem-progenitor cell expansion

We next investigated potential molecular mechanisms whereby elevated integrin-mechanosignaling could expand the stem-progenitor pool of MECs. The mammary gland is a hormonally regulated tissue whose development, differentiation and function are regulated by estrogens, progesterone and prolactin. In particular, menstrual cycle hormone fluctuations influence MEC proliferation and the high progesterone levels present during diestrus can promote MEC progenitor cell expansion^33–35^. Consistently, qRT-PCR analysis for EMT and MEC progenitor cell associated genes of basal and luminal sorted cells isolated from the V737N mice, synchronized to be in vaginal cytology-confirmed diestrus (Fig. 4a)^62^, showed significant increases in *Smarca4*, *Yap1*, *Bmi1*, *Wwtr1 (Taz)*, *Smarca2*, *Ctnnb1*, *Myc*, and *Sox9* (Fig. 4b-h) in the basal MECs as compared to the same cells isolated from CTL mice. By contrast, these differences were not evident in the MECs isolated from mice in estrus (*Wwtr1 (Taz)*, *Snai2*, *Ezh2*, *Bmi1*, *Smarca2*, *Smarca4*, *Myc*, *Sox9*, *Yap1*, and *Ctnnb1;* Extended Data Fig. 7a-i). After confirming no significant differences in transcript levels for the estrogen or progesterone receptors (*Esr1*, *Pgr (A and B)*, *Pgr (B*)) in MECs isolated from V737N and CTL mice (Extended Data Fig. 8a-c), we turned our focus towards examining the impact of paracrine factors known to influence stem-progenitor MEC expansion. Progesterone is one such paracrine factor that stimulates the production of RANKL and WNT4 from luminal Progesterone Receptor (PR)-positive MECs to induce MEC proliferation and progenitor expansion (Fig. 4i)^33,34,63^. To begin with, an ELISA assay detected higher RANKL protein in the mammary glands excised from the V737N mice (Fig. 4j). Therefore, we further analyzed *Tnfsf11* (*Rankl)* and *Wnt4* gene expression in basal and luminal MECs isolated from the V737N and CTL mice. Interestingly, we observed that *Tnfsf11* (*Rankl)* but not *Wnt4* transcript levels was expressed at significantly higher levels in V737N-β1 integrin positive luminal MECs, while *Tnfrsf11a* (*Rank)* transcript levels were slightly increased in both luminal and basal V737N MECs (Fig. 4k,l and Extended Data Fig. 8d). Consistently, cell cycle transcriptional targets of RANKL, namely *Ccnd1* and *Ccnd2*, were upregulated in the V737N-β1 integrin expressing luminal and basal MECs, respectively, when compared to their corresponding isolated CTL MECs (Fig. 4m,n). These results suggest the higher frequency of stem-progenitor cells induced by elevated MEC integrin-mechanosignaling was likely mediated through increased levels of progesterone-stimulated RANKL.

**Fig. 4:**
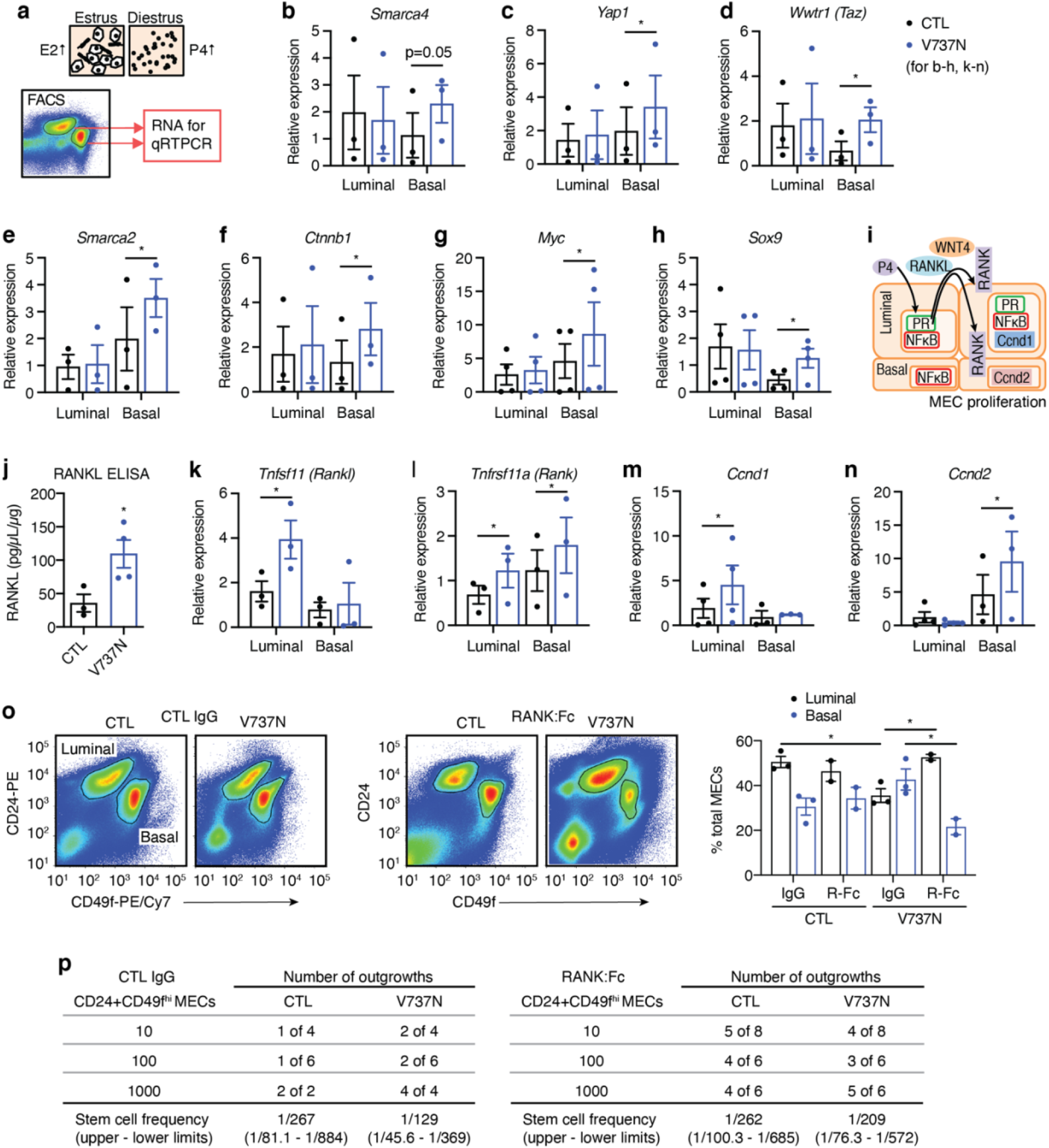
Hormone induced RANKL is critical for mechanosignaling induced mammary progenitor expansion. **a**, Cartoon illustrating cytology used to determine the estrus (estrogen; E2 high) and diestrus (progesterone; P4 high) stages of the estrus cycle in mice prior to FACS-mediated isolation of luminal and basal MECs from CTL and V737N mice for RNA extraction and qRT-PCR. **b-h**, Graphs showing qRT-PCR analysis of relative gene expression for the indicated genes in cells sorted as described in a (n=3-4 independent isolations). **i**, Cartoon depicting progesterone (P4) action in the mammary gland. P4 stimulates progesterone receptor (PR) mediated transcription in luminal epithelial cells. PR-induced paracrine factors, Rankl and Wnt4 stimulate NFκB activity and proliferative responses in LEC and basal cells that include induction of *Ccnd1* and *Ccnd2* transcripts. **j**, Whole mammary glands were lysed and subjected to ELISA to measure RANKL levels (CTL; n=3, V737N; n=4). **k-n**, Graphs showing qRT-PCR analysis of relative gene expression for the indicated genes in MECs sorted as described in a (n=3-4 independent isolations). **o**, MECs were isolated by FACS from CTL and V737N mice and treated with RANK-Fc or IgG control three times per week for two weeks. Representative FACS plots are shown (left; CTL IgG treated, and middle; RANK:Fc treated) and the average percentage of luminal (CD24^+^; CD49f^lo^) and basal (CD24^+^; CD49f^hi^) MECs are plotted (right) (n=2-3 independent mice per group). **p**, Limiting dilution transplantation assays were performed with basal (CD24^+^; CD49f^hi^) MECs sorted from CTL and V737N mice treated as in n (left table; CTL IgG treated, and right table; RANK:Fc treated). Stem cell frequency was calculated from the number of mammary gland repopulating events. All graphs are presented as mean +/- S.E.M. Statistical tests used were paired two-way ANOVA (with two-stage linear step-up procedure of Benjamini, Krieger and Yekutieli for multiple comparisons test) (b-h, k-n), unpaired *t*-test (j) and two-way ANOVA (with Tukey’s multiple comparisons test) (o). *P<0.05, **P<0.005, ***P<0.0005, ns=non-significant.

To causally implicate RANKL in integrin-mechanosignaling induced MEC stem-progenitor expansion, cohorts of the V737N and CTL mice were treated with a RANKL inhibitor (RANK:Fc) over a two week period after which their MECs were collected and the basal cell population was isolated by FACs. The isolated basal MECs were analyzed for their stem-progenitor phenotype by assaying for their ability to repopulate a cleared mammary fat pad using the limited dilution transplantation assay. To begin with FACS analysis of luminal:basal MEC ratios confirmed that RANK:Fc treatment eliminated a large proportion of the basal MEC population in the V737N-β1 integrin expressing mammary glands (Fig. 4o). Moreover, the limiting dilution transplantation assay confirmed that RANK:Fc treatment reduced stem-progenitor frequency in the basal mammary population isolated from V737N mice towards that demonstrated by CTL mice (Fig. 4p). These data imply that integrin mechanosignaling increases RANKL levels in the mammary gland in a progesterone-dependent manner and suggest that this expands the number of stem-progenitor MECs.

### Progesterone receptor activity is augmented by mechanosignaling and matrix stiffness

The progesterone receptor (PR) is phosphorylated post-translationally on several residues to alter its activity (Figure 5A)^64^. ERK-mediated phosphorylation of Serine 294 (S294) is associated with elevated PR transcriptional activity and has been implicated in the specific upregulation of tumor associated stemness-progenitor genes^65–68^. Immunofluorescence staining of mammary gland tissue sections from V737N mice showed greater levels of nuclear PR in luminal MECs, consistent with higher PR and RANKL-signaling (Fig. 5b). These results were confirmed by immunoblot analysis of FACS-isolated luminal mouse MECs, which revealed there was a marked increase in the level of phospho-ERK and phospho-PR (S294) as well as elevated nuclear p65-NFκB levels in the luminal MECs isolated from the V737N mice as compared to those isolated from CTL mouse mammary glands (Fig. 5c,d).

**Fig 5:**
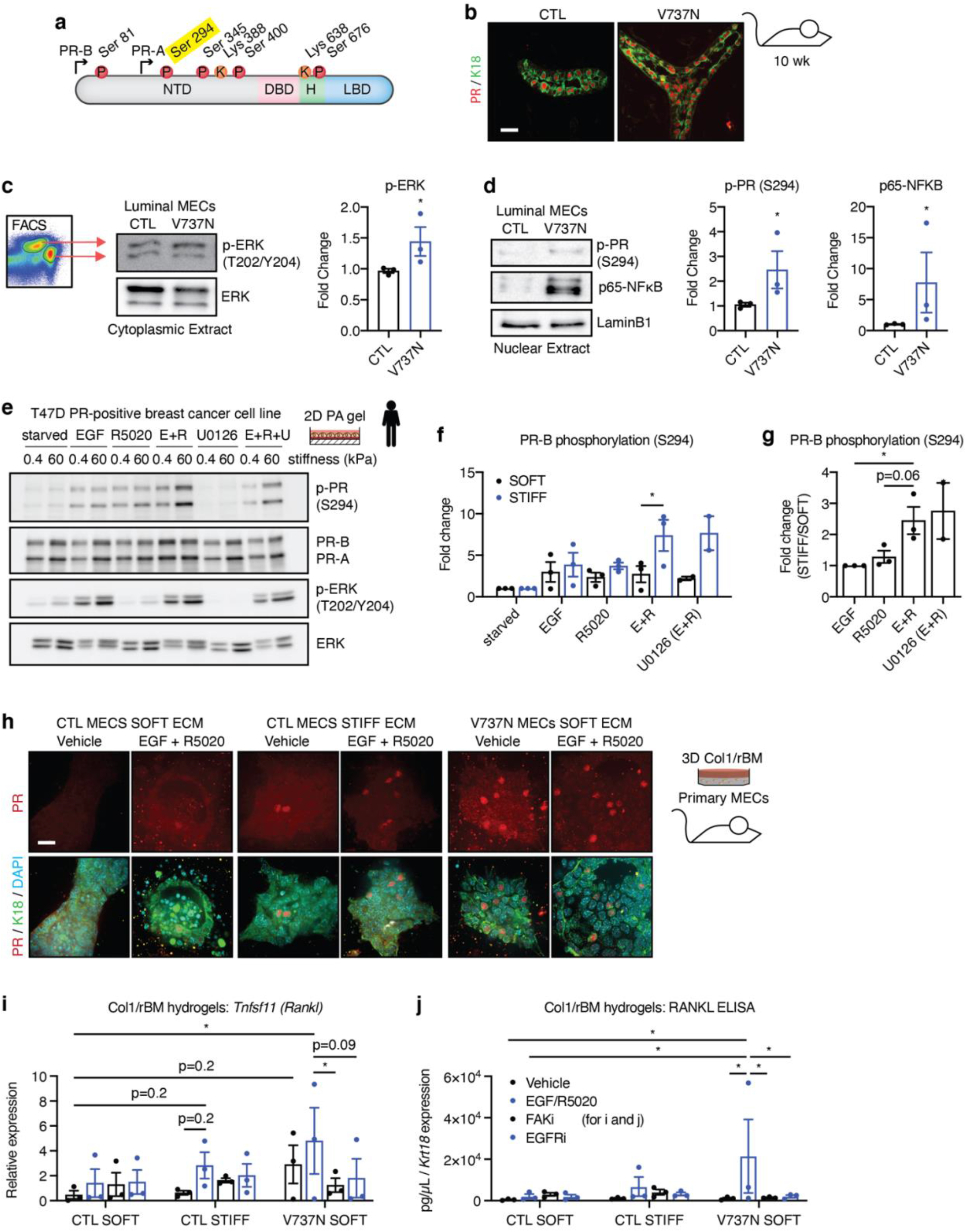
Extracellular matrix stiffness and high mechanosignaling potentiate progesterone signaling. **a**, Cartoon depicting progesterone receptor (PR) isoforms and known sites of post-translational modification (P=phosphorylation, K=SUMOylation/acetylation). **b**, Representative images showing immunofluorescence staining of frozen mammary gland sections from CTL and V737N mice with antibodies specific to progesterone receptor (PR; red) and keratin 18 (K18; green). Scale bar, 50 μm. **c**, Luminal and basal MECs were isolated by FACS as described in Figure 3f. (left) Cells were lysed as nuclear and cytoplasmic fractions prior to immunoblotting with antibodies specific to phospho-ERK (p-ERK; T202/204) and total ERK for the cytoplasmic fraction. (right) Densitometry for phospho-ERK relative to total ERK was calculated for 3 independent cell isolations and plotted. **d**, (left) The nuclear extract of cells isolated as in c were subjected to immunoblot analysis with antibodies specific to phosphorylated progesterone receptor (p-PR, S294), p65-NFκB and Lamin B1 as a loading control. Densitometry for p-PR (S294; middle) and p65-NFκB (right) relative to LaminB1 was measured for 3 independent cell isolations and plotted. **e**, T47D breast cancer cells were cultured on fibronectin conjugated polyacrylamide gels of varying stiffness (0.4 and 6kPa), serum starved overnight, and treated with EGF (20ng/mL), R5020 (10nM), EGF and R5020 together (E+R), U0126 (MEK inhibitor), or U0126 for 1hr prior to EGF and R5020 (E+R+U) for 15 min. Cells were then lysed for immunoblotting with antibodies specific to phospho-PR (S294), PR-A and B, phospho-ERK (p-ERK; T202/204) and ERK. **f**, Bar graph showing densitometry measurements for PR-B phosphorylation relative to PR-B total levels from three independent experiments conducted as in f (SOFT=0.4kPa; STIFF=6kPa). Treatment conditions were normalized to the non-treated serum-starved condition. **g**, Bar graph of the data from f represented as fold change of STIFF/SOFT for each treatment condition. **h**, Primary mouse MECs were isolated from CTL and V737N mice and cultured as organoids in SOFT (no L-ribose) and STIFF (crosslinked with L-ribose) Col1/rBM hydrogels for 24 hr with serum starvation overnight prior to stimulation with vehicle or EGF and R5020 for 15 min. Nuclear translocation of PR was assessed by immunofluorescence staining with antibodies specific for the progesterone receptor (PR, red) and keratin 18 (green), and nuclei (cyan) were stained with DAPI. Representative images are shown. Scale bar, 25 μm. **i**, Primary mouse MECs prepared and cultured as in i were serum starved overnight prior to stimulation with vehicle or EGF/R5020 for 48 hr. Some samples were also treated with FAK-inhibitor (FAK-i, PND1186) or EGFR-inhibitor (EGFRi, Tyrphostin/AG-1478) prior to collection of conditioned media and RNA extraction for qRT-PCR analysis of relative *Tnsfs11* (*Rankl)* gene expression levels (n=3 independent isolations). **j**, The conditioned media from j was subjected to ELISA for determination of soluble RANKL levels (n=3 independent isolations). All graphs are presented as mean +/- S.E.M. Statistical tests used were Mann-Whitney test (c and d, one-tailed) and paired two-way ANOVA (with two-stage linear step-up procedure of Benjamini, Krieger and Yekutieli for multiple comparisons test) (f, g, i, j). *P<0.05, **P<0.005, ***P<0.0005, ns=non-significant.

We next explored whether ECM stiffness could similarly enhance PR-dependent *Rankl* transcription and if this reflected enhanced PR phosphorylation at S294. We cultured PR expressing T47D and MCF7 breast cancer cells on soft (400 Pascals) and stiff (60 kPa) fibronectin coated polyacrylamide gels and monitored for effects on PR phosphorylation in response to treatment with epidermal growth factor (EGF) and the synthetic progestogen/PR-agonist (promegestone/R5020) either in combination or as single agents. Immunoblots of whole cell lysates isolated from T47D and MCF7 breast cancer cells revealed that a stiff ECM synergistically enhanced the ability of EGF and R5020 to stimulate S294 phosphorylation of PR (2-fold induction; Fig. 5e-g; Extended Data Fig. 9a-c).

To verify whether a stiff ECM could promote PR-dependent transcription of *Tnfsf11* (*Rankl)* we isolated mammary organoids from CTL and V737N mice. We then cultured these organoids within SOFT and STIFF Col1/rBM ECM hydrogels and assayed for PR nuclear translocation and *Tnfsf11* (*Rankl)* expression following treatment with EGF and R5020^25,48,55^. Immunofluorescence staining demonstrated significantly greater nuclear translocation of PR in the EGF/R5020-treated organoids embedded within the STIFF Col1/rBM gels as well as in V737N organoids within the SOFT Col1/rBM gels regardless of treatment (Fig. 5h). qRT-PCR analysis revealed that CTL organoids within the STIFF Col1/rBM gels and the V737N β1 integrin expressing organoids within the SOFT Col1/rBM gels also expressed higher levels of *Tnfsf11* (*Rankl)* versus CTL organoids in SOFT Col1/rBM gels, and an ELISA of conditioned media from these cultures confirmed elevated protein levels of RANKL (Fig. 5i,j). Treatment with a FAKi or an Epidermal Growth Factor Receptor (EGFR) inhibitor (EGFRi; Tyrphostin/AG1478) abrogated ECM stiffness and mechanosignaling induced RANKL levels (Fig. 5i,j). These data indicate that a stiff matrix and elevated integrin-mechanosignaling potentiate progesterone-induced RANKL signaling in MECs, at least in part by enhancing ERK-mediated phosphorylation of PR.

### Stiff tumors and tissues with high mammographic density have elevated RANKL pathway and mammary epithelial progenitor activity

Based on our findings that integrin-mechanosignaling potentiates progesterone-dependent RANKL expression to expand the stem-progenitor MEC pool, we examined whether the pro-tumorigenic, EMT-stem-progenitor phenotype we documented to be induced by a stiffened ECM and elevated integrin-mechanosignaling was accompanied by elevated RANK signaling in mammary tumor cells. In agreement with this paradigm, qRT-PCR analysis and immunofluorescence staining of the HER2-positive PDX tumors that were grown in immunocompromised mice within the STIFF Col1/rBM ECM stroma had high RANKL transcript and protein levels when compared to tumors grown within the SOFT Col1/rBM ECM stroma (Fig. 6a-c). By contrast, although *WNT4* transcript levels were increased modestly the levels were not significantly higher in these PDX tumors (Extended Data Fig. 10a). Moreover, although we failed to detect any increase in *TNFRSF11A* (*RANK)* transcript in these tumors, immunofluorescence staining indicated that those syngeneic tumors that developed within the STIFF ECM stroma had significantly elevated RANK protein and higher RANK signaling as indicated by a strong presence of nuclear p65-NFκB levels (Extended Data Fig. 10b-d). Consistently, syngeneic PyMT tumors embedded within STIFF ECM stroma and the V737N^Neu^ tumors with elevated integrin mechanosignaling also expressed higher levels of *Tnsfs11* (*Rankl)* mRNA and RANKL protein and exhibited strong staining for RANK receptor and nuclear p65-NFkB as compared to their respective control mammary tumors (Extended Data Fig. 10e-p). These findings are consistent with tissue tension enhancing RANK signaling to promote tumor aggression by expanding the stem-progenitor population.

**Fig. 6:**
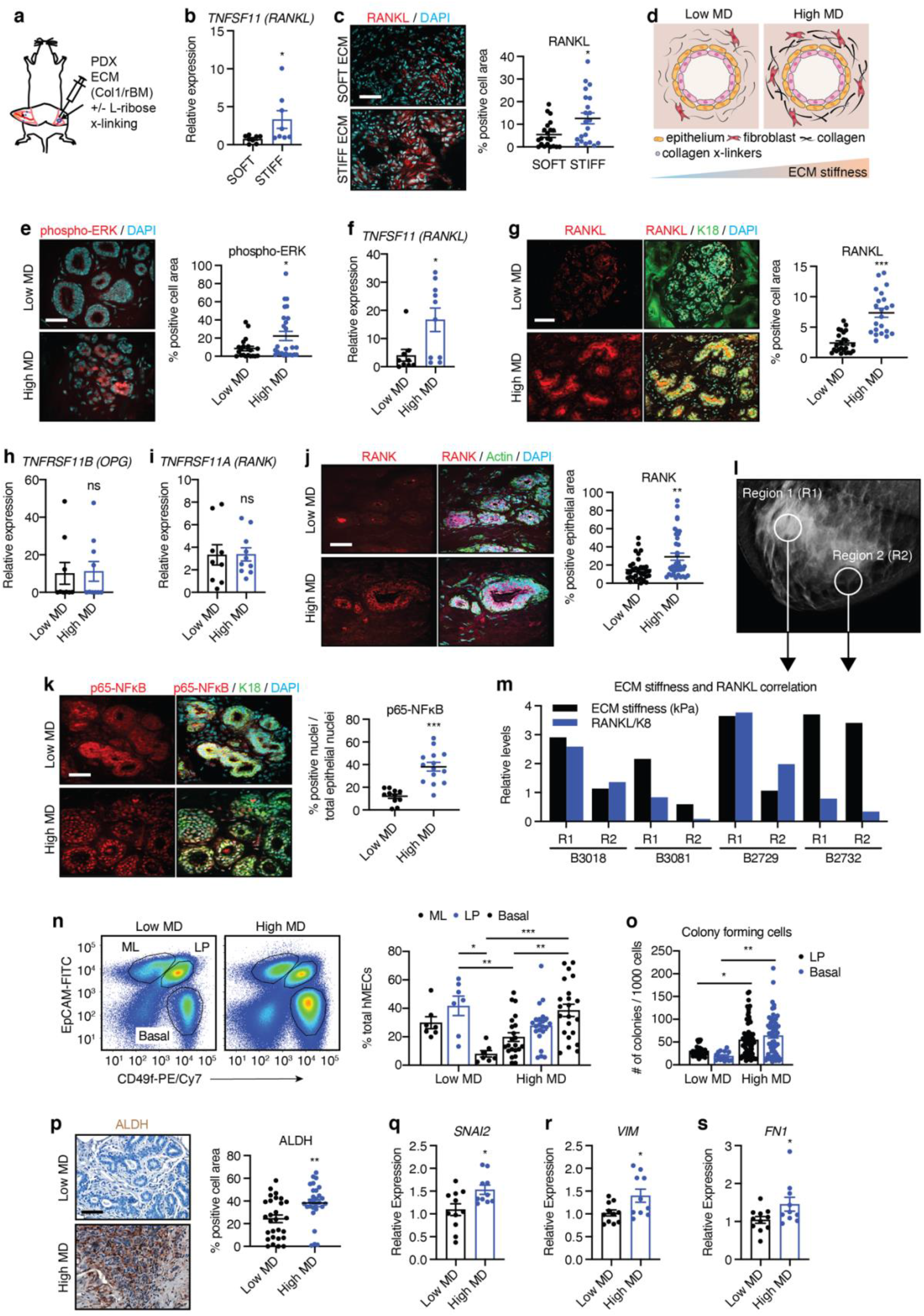
Extracellular matrix stiffness is associated with elevated RANK activity and breast cancer risk. **a**, Cartoon of HER2-positive patient derived xenografts (PDXs) with Col1/rBM ECM stroma implantation strategy with (STIFF) and without (SOFT) L-ribose mediated crosslinking (x-linking). **b**, qRT-PCR analysis of relative gene expression for *TNFSF11* (*RANKL)* in HER2-positive PDX tumors developed in SOFT (n=8) and STIFF (n=8) stromal matrices. **c**, (left) Representative images of immunofluorescence staining for frozen tumor sections as in b with an antibody specific to RANKL (red). Nuclei were counterstained with DAPI (cyan) (n=6 for SOFT and STIFF). Scale bar, 50 μm. (right) Graph showing average percentage of RANKL positive staining per total cell area. **d**, Cartoon depicting human breast tissue organization with low and high mammographic density (MD). High MD tissues have more abundant epithelium, fibroblasts, collagen matrix and collagen crosslinking enzymes contributing to a stiffer stroma. **e**, (left) Representative images of immunofluorescence staining for frozen human breast tissue sections (n=6 for low and high MD) with an antibody specific for phosphorylated-ERK (Red). Nuclei are counterstained with DAPI (cyan). Scale bar, 50 μm. (right) Graph showing the average percentage of positive phospho-ERK staining per total cell area. **f**, qRT-PCR analysis of relative gene expression for *TNFSF11 (RANKL)* (normalized to *Krt8*) in low and high MD human breast tissues (n=9; low MD, n=10, high MD). **g**, (left) Representative images of immunofluorescence staining for frozen human breast tissue sections (n=6 for low and high MD) with antibodies specific for RANKL (Red) and Keratin 18 (K18, green). Nuclei are stained with DAPI (cyan). Scale bar, 50 μm. (right) Graph showing average percentage of positive RANKL staining per total cell area. **h-i**, qRT-PCR analysis as in f for *TNFRS11B (OPG)* and *TNFRSF11A (RANK)* (n=9-10 for low and high MD). **j**, (left) Representative images of immunofluorescence staining for frozen human breast tissue sections (n=6 for Low and high MD) with an antibody specific for RANK (Red). Actin filaments are stained with Phalloidin (488, green) and nuclei are counterstained with DAPI (cyan). Scale bar, 50 μm. (right) Graph showing average percentage of positive RANK staining per total cell area. **k**, (left) Representative images of immunofluorescence staining for human breast tissue frozen sections (n=8 for low and high MD) with antibodies specific for p65-NFκB (Red) and Keratin 18 (K18, green). Nuclei are counterstained with DAPI (cyan). Scale bar, 50 μm. (right) Graph showing average percentage of positive nuclear p65-NFκB staining per total cell nuclei. **l**, Example of a whole breast mammogram where two regions of different mammographic density (MD) were excised for storage as OCT-embedded frozen tissue blocks. **m**, Frozen sections from two regions of the same breast as depicted in l were used to measure average ECM stiffness by AFM. Sections of each region were also used to isolate RNA and determine relative *TNFSF11* (*RANKL)* gene expression as in f. Average AFM measurements and relative *RANKL* gene expression levels were plotted for each region to display their relationship. B numbers represent individual patients (n=4). **n**, (left) Representative plots showing FACS analysis of low MD (n=7) and high MD (n=22) human breast tissues using the epithelial surface antigens, EpCAM and CD49f. (right) Graph illustrating the average percentage of mature luminal (ML, EpCAM^hi^; CD49f^lo^), luminal progenitor (LP, EpCAM^hi^; CD49f^hi^) and basal (ML, EpCAM^lo^; CD49f^hi^) MECs. **o**, FACS isolated (as in r, n=7; low MD, n=22, high MD) LP and Basal MECs were subjected to colony formation assays in rBM to assess progenitor activity. Graph shows the average number of colonies per 1000 cells formed for both lineages from low and high MD tissues. **p**, (left) Representative images of Immunohistochemical staining of paraffin sections from low MD (n=7) and high MD (n=6) breast tissues with an antibody specific for ALDH. Nuclei are counterstained with hematoxylin. Scale bar, 50 μm. (right) Graph showing average percent positive ALDH staining per total cell area. **q-s**, Graphs showing qRT-PCR analysis of relative gene expression (normalized to *Krt8*) for the indicated genes in low MD (n=11) and high MD (n=10) human breast tissues. All graphs are presented as mean +/- S.E.M. Statistical tests used were unpaired *t*-test (b, f, h, i, q-s), Mann-Whitney test (c, e, g, j, k, p) and two-way ANOVA (with Tukey’s multiple comparisons test) (n, o). *P<0.03, **P<0.002, ***P<0.0002, ns=non-significant.

Women with high mammographic density (high MD), have a four-fold increased lifetime risk to malignancy and their breast tissue contains more fibroglandular tissue and higher levels of epithelial proliferation than the breast tissue classified as low MD^48,69–71^. We recently showed that the stroma from tissues with high MD not only has more abundant fibrillar interstitial collagen and stromal fibroblasts that express higher levels of the collagen cross-linking enzymes lysyl oxidase (LOX) and lysyl hydroxylase 2 (LH2), but that their stroma is significantly stiffer than that surrounding the ductal-lobular epithelium in low MD tissues (Figure 6D)^48^. Given prior studies linking stem-progenitor frequency and cellular proliferation levels to cancer risk^1–3^ we examined the stiff, high MD tissue from premenopausal women for evidence of elevated RANK signaling and a higher frequency of stem-progenitor MECs. To begin with, we noted that the epithelium of high MD tissue had higher levels of phospho-ERK compared to the epithelium in the low MD breast tissue (Fig. 6e). Furthermore, qPCR revealed elevated *TNSFS11 (RANKL)* levels in the high MD tissue that immunofluorescence staining confirmed was translated to higher RANKL protein levels (Fig. 6f,g). Importantly, however, we did not detect any differences in expression at the mRNA level for either the RANKL decoy receptor, *TNFRSF11B (OPG)* or *TNFRSF11A (RANK)* in the high and low MD tissues (Fig. 6h,i). Consistent with the mammary tumor tissues analyzed, we did observe an increase in RANK protein expression by immunofluorescence (Fig. 6j). Likewise, we quantified higher levels of nuclear p65-NFkB indicating greater RANKL signaling activity (Fig. 6k). To reduce potential variability linked to systemic and tissue levels of progesterone, we next compared the RANKL levels from two regions of the same breast reflecting high and low MD with AFM-determined stromal stiffness (Fig. 6l). Once again, we observed a strong correlation between *TNSFS11* (*RANKL)* expression and stromal stiffness for each patient set (2 regions) from 4 individual patients (Fig. 6m). These findings suggested that the documented elevated breast cancer risk in women with high MD may at least in part, be linked to an increase in the frequency of stem-progenitor cells.

To investigate whether the breast tissue from women with high MD contained a higher number of stem-progenitor cells we utilized breast tissue surgically excised from prophylactic mastectomy and classified as high and low MD and performed FACS analysis of the MEC populations using EpCAM and CD49f to distinguish mature luminal (ML), from luminal progenitor (LP) and Basal MECs (Fig. 6n). We observed a striking and highly significant increase in the relative abundance of the basal MEC compartment in the tissues classified as having higher MD (Fig. 6n). The stem-progenitor phenotype of the MECs isolated from the high MD breasts was functionally confirmed by showing higher colony formation capacity over the breast epithelium isolated from tissues with low MD (Fig. 6o). IHC staining for ALDH, which is associated with human mammary progenitor epithelial cells, was also increased in the breast tissue isolated from women with high MD (Fig. 6p). Given prior studies and our results in experimental models linking a mesenchymal phenotype to epithelial stem-progenitor activity we next assayed the high and low MD tissues for EMT markers. Not surprisingly, we noted that the mRNA for several mesenchymal markers including *SNAI1*, *VIM* and *FN1* were expressed at higher levels in the breast tissues classified as having high MD as compared to those with low MD (Fig. 6q-s). These data imply that women with high MD may be predisposed to developing breast cancer, not only because they have a higher number of MECs that proliferate more, but that they also contain a higher frequency of tissue resident long-lived stem-progenitor cells. Our findings further suggest the elevated stem-progenitor MEC frequency we documented in breast tumors and in the normal breast with high MD may be due to stiffness-enhanced progesterone-induction of RANKL and elevated RANK signaling.

## DISCUSSION

Here we demonstrate, for the first time, that a stiff stroma expands the resident population of mammary epithelial stem-progenitor cells. Our findings highlight the critical role played by physical cues in regulating stem-progenitor cell behavior and are consistent with prior data implicating tissue mechanics in stem-progenitor cell proliferation and viability^12,14,15,72–75^. We elaborate upon these prior studies by identifying a molecular mechanism whereby ECM stiffness increases stem-progenitor cell frequency through the tension-dependent enhancement of β1 integrin mechanosignaling that potentiates progesterone induced RANK activity. Accordingly, our results provide one plausible explanation for why deleting β1 integrin in the MEC basal population so dramatically reduces basal progenitor activity in the tissue to favor luminal differentiation^76^, and why deleting FAK in the entire breast epithelium so potently impairs luminal and basal progenitor cell function^77^. Our findings could also explain links between β1 integrin signaling and FAK activity and breast tumor aggression and metastasis^78–87^. Indeed, given that a basal-like luminal progenitor cell was recently identified and that HER2-positive and basal-like human breast cancers are thought to originate from luminal progenitor MEC populations^88,89^, this raises the possibility that β1 integrin mechanosignaling favors the emergence of aggressive mesenchymal-like breast cancer subtypes by expanding a pool of basal-like progenitor MECs.

Conceptually, our finding that a stiff ECM potentiates progesterone signaling is consistent with earlier work illustrating how a stiff ECM potentiates cell growth and survival by enhancing the context of growth factor receptor and GPCR signaling and expands this paradigm to include modification of hormonal signaling and transcription^14,19,24–31^. This raises the intriguing possibility that tissue tension compromises tissue homeostasis and promotes cancer aggression in other tissues by regulating responses to other tissue-specific hormones including growth hormone/IGF-1, testosterone and ghrelin^90–93^. Importantly, to our knowledge, this is the first evidence demonstrating that ECM stiffness and mechanosignaling are critical modulators of PR signaling, progenitor-like MEC function and a mesenchymal breast cancer phenotype. These data are consistent with reports that ECM remodeling through ovarian hormone induced ADAMTS18 is important for the maintenance of mammary progenitor cells through hippo and Wnt signaling^94^. Moreover, we previously showed that V737N-mediated mechanosignaling disrupts membrane β-catenin localization in MECs^26^, which is indicative of active Wnt signaling and in agreement with observations that PR-induced RANKL/WNT4 can foster early dissemination or HER2-induced mammary tumor cells^95^. In the present study we have mechanistically linked stromal stiffness and mechanosignaling to hormone induced progenitor cell expansion by demonstrating that these factors potentiate ERK mediated PR phosphorylation at S294, which has also been associated with enhanced cancer stem cell associated gene expression in breast cancer^67^. Interestingly, PR phosphorylation at S294 leads to enhanced degradation of PR, thereby hindering its detection by immunohistochemistry, which may explain potential contradiction between these data and reports of progesterone activity as being beneficial for breast cancer prognosis^67,96^. Our present data suggest that interventions aimed at reducing stromal stiffness or blocking the proposed PR-sensitive mechanistic pathway may prove useful for preventing tumor initiation and aggression.

We quantified higher RANKL and RANK levels in the breast tumors that developed within a highly fibrotic, stiff ECM stroma, as well as in the normal human breast tissue with high MD and a stiffer stroma^48^. We further demonstrated that tumors developing within a highly fibrotic, stiffened ECM stroma have a significantly higher frequency of stem-progenitor like tumor cells and that the normal tissue in women with high MD have an increased frequency of functional mammary epithelial basal-like progenitor cells. These findings are consistent with prior data showing that tissues with BRCA1 mutation possess high RANK expression and harbor a higher frequency of progenitor-like cells, as well as reports that more aggressive subtypes of breast cancer that typically develop a more profound fibrosis also have elevated RANK expression^97–99^. Indeed, and consistent with our findings, RANK signaling has been previously demonstrated to promote an EMT in normal MECs and in breast cancer cells^100,101^. Our data expand these findings to suggest that many of these effects that contribute to greater risk of tumor initiation and promote the evolution of aggressive cancers, may be mediated by an underlying stiff stroma and elevated MEC mechanosignaling. Not only did we observe more RANKL and RANK in high MD tissues and mammary tumors, but we also observed elevated levels of nuclear p65-NFκB indicating not only elevated RANK signaling but also potentially more cytokine expression. Such observations suggest that a stiff ECM stroma might additionally promote breast cancer risk and drive tumor aggression by stimulating tissue inflammation and by altering anti-tumor immunity. Indeed, women with high MD have more infiltrating myeloid cells and more fibrotic TNBCs that possess a stiffer stroma also have the highest density of infiltrating inflammatory myeloid cells^17,102^. As such, these data provide a plausible explanation for the link between tissue fibrosis and inflammation, as they imply there exists a positive reinforcing loop whereby fibrosis stiffens the ECM, which in turn potentiates NFκB activity to stimulate expression of pro-inflammatory cytokines that thereafter recruit the infiltration of pro inflammatory myeloid cells into the tissue. This proposed mechanism would not only explain how tissue fibrosis and stiffness drive breast cancer progression and metastasis but could also explain the higher risk to malignancy observed in women with high MD as well as those with a BRCA1-mutation^97,103–106^. Interestingly, several immune populations produce and respond to RANKL^107^, and FoxP3-positive regulatory T cell (Treg)-derived RANKL promote RANK-positive tumor cell metastasis in breast cancers^108^. Treg cells may be recruited by αSMA-positive stromal fibroblasts that are an abundant cellular constituent in fibrotic, stiff tumors. Thus, the stiffened, fibrotic tumor stroma could enhance the recruitment of Treg cells that thereafter drive tumor aggression and stem-progenitor cell expansion by secreting RANKL and enhancing the recruitment of myeloid cells through NFκB activation and elevated cytokine expression. Indeed, RANKL inhibition limits tumor progression by improving CD8-positive T cell responses alone and in combination with immune checkpoint inhibitors^109–111^. Our findings thus provide new evidence supporting the use of RANKL inhibition as an effective anti-cancer treatment and as a prevention modality, particularly in individuals with known or measured high ECM stiffness and activated integrin-mechanosignaling, such as those with high MD.

## METHODS

### RNA-seq library preparation, sequencing, and analysis

RNA was isolated using TRIzol (Invitrogen, Cat. #: 15596018) followed by chloroform extraction. RNAseq library preparation was performed by the Functional Genomics Laboratory (FGL), a QB3-Berkeley Core Research Facility at UC Berkeley. Total RNA samples were checked on a Bioanalyzer (Agilent) for quality and only high-quality RNA samples (RIN > 8) were used. At the FGL, Oligo (dT)25 magnetic beads (Thermofisher) were used to enrich mRNA, and the treated RNAs were rechecked on the Bioanalyzer for their integrity. The library preparation for sequencing was done on Biomek FX (Beckman) with the KAPA hyper prep kit for RNA (now Roche). Truncated universal stub adapters were used for ligation, and indexed primers were used during PCR amplification to complete the adapters and to enrich the libraries for adapter-ligated fragments. Samples were checked for quality on an AATI (now Agilent) Fragment Analyzer. Samples were then transferred to the Vincent J. Coates Genomics Sequencing Laboratory (GSL), another QB3-Berkeley Core Research Facility at UC Berkeley, where Illumina sequencing libraries were prepared. qPCR was used to calculate sequence-able molarity with the KAPA Biosystems Illumina Quant qPCR Kits on a BioRad CFX Connect thermal cycler. Libraries were pooled evenly by molarity and sequenced on an Illumina NovaSeq6000 150PE S4. Raw sequencing data were converted into fastq format, sample-specific files using the Illumina bcl2fastq2 software on the sequencing centers local Linux server system. RNAseq fastq files were mapped to the primary assembly of the Gencode v33 human genome using Rsubread (version 2.0.1) and counted using featureCounts. Lowly expressed genes were filtered out if they did not have at least one CPM in at least 4 samples. Data normalization was performed using calcNormFactors in edgeR (version 3.28.1). Gene ontology was performed using Gage (version 2.36.0) with gene lists from MsigDB version 7.2.

### Human Breast Tissue Collection and Processing

Tissue specimens were collected from prophylactic mastectomy and 2×2 cm pieces were formalin-fixed and paraffin embedded (FFPE), or flash frozen in OCT (Tissue-Tek) by slow immersion in liquid nitrogen or placement on dry ice. Similarly sized fragments were also collected as fresh tissue with immersion in media (phenol red free-DMEM/F12) with 10% charcoal stripped fetal bovine serum (FBS Benchmark, Cat. #: 100-106) and GlutaMAX (Gibco, Cat. #: 35050-061) supplementation for transportation to the Weaver laboratory at UCSF. Fresh tissue was used for FACS-mediated human MEC isolation and subsequent culture in colony formation assays or for RNA extraction. PDX tissues were obtained from Dr. Alana Welm (Huntsman Cancer Institute, University of Utah) and Dr. Michael Lewis (Baylor College of Medicine)^57,58^. All human breast tissue specimens were collected prospectively from consenting patients (informed consent provided prior to surgery) undergoing surgery at the University of California, San Francisco, (UCSF) or Duke University Medical Center between 2010 and 2020. Samples were stored and analyzed with deidentified labels to protect patient data in accordance with the procedures outlined in the Institutional Review Board Protocol #10-03832, approved by the UCSF Committee of Human Resources and the Duke University IRB (Pro00054515).

### Animals and Animal Care

Animal husbandry and all procedures on mice were carried out in Laboratory Animal Resource Center (LARC) facilities at UCSF Parnassus in accordance with the guidelines stipulated by the Institutional Animal Care Use Committee (IACUC) protocols, #AN133001 and #AN179766, which adhere to the NIH Guide for the Care and Use of Laboratory Animals. FVB/NJ, C57Bl6/J, and NOD/SCID mice were purchased from Jackson Laboratories for orthotopic implantation assays. LSL-V737N-β1 transgenic mice were generated as described^26^ and maintained on an FVB/NJ background. MMTV-Cre and MMTV-Neu mice were purchased from Jackson laboratories on an FvBN/J background and crossed with LSL-V737N-β1 mice. For lineage tracing, K5-Cre/ERT2 mice were obtained from Dr. Ophir Klein (UCSF) and MMTV-PyMT/R26-Confetti mice^112^ were obtained from Dr. Mikala Egeblad (Cold Spring Harbor) and maintained on a C57Bl6/NJ background. MMTV-PyMT tumor cells were derived from mice on a C57Bl6/NJ background and were implanted into syngeneic mice for tumorigenesis assays.

### Lineage tracing in mice

Lineage tracing in mice was performed using R26-Confetti reporter mice, which were bred with LSL-V737N-β1 mice and K5-Cre/ERT2 mice. Lineage tracing and V737N-β1 integrin expression were induced primarily in basal lineage MECs by a single intraperitoneal injection of Tamoxifen (Cayman Chemical, Cat. #: 13258, 1.5 mg) when mice were 3-4 weeks of age. Lineage tracing was assessed at two time points (2 and 16 weeks) post tamoxifen induction, when mammary glands were excised and fixed with 4% paraformaldehyde for 30 min prior to embedding and freezing in OCT. A cryostat was used to cut 40 μm thick mammary gland sections which were then stained with Alexa Fluor (AF) 647 Phalloidin (Invitrogen, Cat. #: A22287) to visualize actin filaments when analyzing lineage clonality using a Leica TCS SP5 (five channels) Confocal Microscope with Leica Application Suite (LAS) software. Clonality was assessed by counting the number of clones present in uniformly sized segments of mammary ductal epithelium and counting the number of cells constituting each clone.

### RANKL inhibition in mice

10-week-old wild type mice and mice expressing V737N-β1 integrin (V737N) were treated with PBS/IgG control antibody or RANK-Fc^113^ (AMGEN, 10 mg/kg) by intraperitoneal injection 3 times per week for a period of two weeks (6 total injections) prior to mammary gland harvest and MEC isolation for limiting dilution transplantation assays.

### Collagen/rBM hyrdrogels with orthotopic implantation of tumor cells

Rat tail collagen-1 (High concentration, Corning, Cat. #: 354249) was incubated with 0.1% acetic acid (non-crosslinked; SOFT) or 0.1% acetic acid with 500 mM L-ribose (Chem Impex International, Cat. #: 28127) (cross-linked; STIFF) for at least 10 days before preparation of Col1/ rBM hydrogels for orthotopic implantation of tumor cells or tumor fragments^26,48^. Col1 mixtures were then combined with basement membrane extract (R&D Systems, Cultrex BME, type 2, Pathclear, Cat. #: 3532-005-02) (20% final volume), PBS and 1N NaOH to a slightly acidic pH (pH ∼6.5) as determined by pH strips. PyMT tumor cells were resuspended in Col1/rBM preparations and maintained at 4 °C until implantation into the inguinal mammary fat pad of syngeneic mice. For PDX tissues, Col1/rBM with and without L-ribose was injected orthotopically into a cleared inguinal fat pad and allowed to set for 3-5 minutes prior to implantation of a PDX tissue fragment approximately 2×2 mm in size.

### Collagen/rBM hydrogels for primary MEC culture, hormone and inhibitor treatment

Primary mouse MEC organoids were prepared by manual chopping of mouse mammary gland tissues harvested from 10–12-week-old mice followed by digestion with shaking for 1 hr at 37 °C with Collagenase A from Clostridium histolyticum (10 mg/mL; Roche, Cat. #: 13560925) and Hyaluronidase from bovine testes (1 mg/mL, Sigma-Aldrich, Cat. #: H3506) in DMEM/F12 supplemented with 1% FBS. Digests were then pelleted by centrifugation (1200 rpm), washed in PBS with 2% FBS (Wash buffer) and digested for a further 5 min with Dispase II from *Bacillus polymyxa* (2.5 mg/mL, Sigma-Aldrich, Cat. #: D4693) and DNAse I from bovine pancreas (100 μg/mL; Roche, Cat. #: 10104159001). A final digestion with 0.25% Trypsin-EDTA solution was performed for single cell dissociation. Single cells were cultured in rBM (6%) for 10-14 days prior to their recovery from rBM and resuspension in Col1/rBM hydrogels prepared as described^25,48,55^. Hydrogels were prepared from Rat Tail collagen I (Corning) incubated for >10 days with 0.1% acetic acid containing 500mM L-Ribose (Chem Impex International) (crosslinked; STIFF) or 0.1% acetic acid alone (non-crosslnked; SOFT). Collagen was then mixed with 20% rBM, DMEM, PBS and 1 μg/mL Human Plasma Fibronectin Purified Protein (Sigma-Aldrich, Cat. #: FC010). 1N NaOH was added to achieve a neutral ∼pH and a thin base layer of 100 μL volume was added to the well of a 48-well tissue culture plate precoated with 1% agarose. Mammary organoids were resuspended in the SOFT and STIFF collagen preparations and plated as a top layer of 100 μL and allowed to solidify for 30 min at room temperature followed by 30 min at 37 °C. Cell medium was then added, and gels were detached from the sides of the wells to remain suspended in cell medium. Alternatively, following light trypsinization, organoids were immediately cultured in Col1/rBM hydrogels without preparatory culture in 100% rBM. Organoids were cultured in DMEM/F12 supplemented with 20 ng/mL Epidermal Growth Factor (EGF), 10 μg/mL Insulin and 2 μg/mL hydrocortisone. Organoid/hydrogel cultures were serum starved overnight before treatment with EGF (20 ng/mL) and promegestone/R5020 (1, 10, 100 nM; Perkin Elmer, Cat. #: NLP00400) and 10 nM R5020 with or without the addition of FAK (1 μM; PND-1186, Chem Scene, Cat. #: 1061353-68-1) and EGFRi (Tyrphostin/AG1478, Selleck Chem, Cat. #: S2728) inhibitors (FAKi and EGFRi). TRIZol (Invitrogen) was used for RNA extraction using the Ambion mirVana kit (Invitrogen, Cat. #: AM1560) per manufacturer’s instructions.

### Monitoring of Tumor growth and metastasis

Tumor growth was monitored by palpation and caliper measurement weekly or biweekly. Lung metastases were quantified by counting of surface lesions at time of animal sacrifice, and by examination of histological lung sections stained by H&E. Lungs were scanned using a ZEISS Axio Scan.Z1 digital slide scanner equipped with CMOS and color cameras, 10x, 20x and 40x objectives, and lesion area was determined by tracing metastatic lesions in QuPath^114^.

### Quantitative Reverse Transcriptase-polymerase chain reaction (qRT-PCR)

RNA was prepared from FACS-isolated MECs or flash frozen and pulverized mammary tumor tissues using TRIZol reagent (Invitrogen). Reverse transcription reactions were performed using M-MLV reverse transcriptase (Biochain, Cat. #: Z5040002) with random hexamer primers. cDNA was mixed with PerfeCTa SYBR Green FastMix (Quantibio, Cat. #: 95072-05K) for qPCR analysis using an Eppendorf realplex2 epgradient S mastercycler. Thermal cycling conditions were 10 min at 95 °C, followed by 40 cycles of 15s at 95 °C and 45 s at 65 °C. Melting curve analysis was used to verify primer pair specificity. Relative mRNA expression was determined by the ΔΔCT method with normalization to *GAPDH*, *18S* or *KRT8*.

### Quantitative polymerase chain reaction (qPCR) Arrays

Human EMT qPCR arrays were purchased from Qiagen (Cat. #: PAHS-021Z), performed as described using RNA from PDX mammary tumors grown in SOFT and STIFF Col1/rBM hydrogels, and analyzed using available product resources from Qiagen. Selected genes were plotted for presentation in Figure 2 and Supplemental Figure 3.

### Immunohistochemistry (IHC)

IHC was performed as described^51^ using antibodies specific to E-cadherin (BD Biosciences, clone 36, Cat. #: 610181, 1:400) and ALDH (BD Biosciences, clone 44, Cat. #: 611194, 1:200). Briefly, antigen retrieval was accomplished by boiling sections in 10 mM citrate buffer (10min). Following primary antibody incubation overnight at 4 °C, sections were incubated for 1 hr with species-specific Horseradish Peroxidase (HRP)-conjugated secondary antibodies (ImmPRESS HRP Goat Anti-Mouse or Rabbit IgG Polymer Detection Kit, Peroxidase, Vector Laboratories, Cat. #: MP-7452 and MP-7451) before developing positive staining with ImmPACT DAB Substrate Peroxidase (HRP, Vector Laboratories, Cat. #: SK-4105). Images of stained sections were acquired with an Olympus microscope (IX81) and 10x, 20x or 40x objectives.

### Immunofluorescence

Immunofluorescence was performed using the following specific antibodies: phospho-p130-Cas (Y410) (Cell Signaling Technology, Cat. #: 4011, 1:200), phospho-FAK (Y397) (Cell Signaling Technology, Cat. #: 8556, 1:200), phospho-FAK (Y397) (Invitrogen, clone 141-9, Cat. #: 44-625G), phospho-p44/42 MAPK (ERK1/2) (T202/Y204) (Cell Signaling Technology, Cat. #: 9101, 1:200), NFκB p65 (D14E12) XP (Cell Signaling Technology, Cat. #: 8242), phospho-AKT substrates (RXXS*/T*) (110B7E) (Cell Signaling Technology, Cat. #: 9614, 1:400), phospho-Histone H3 (S10) (Cell Signaling Technology, Cat. #: 9701, 1:200), cleaved Caspase-3 (N175) (Cell Signaling Technology, Cat. #: 9661, 1:200), Integrin β1, activated (Sigma-Aldrich, clone HUTS-4, Cat. #: MAB2079Z, 1:400), Integrin β1 (Sigma-Aldrich, clone JB1A, Cat. #: MAB1965, 1:400), PR (Santa Cruz Biotechnology, clone H-190, Cat. #: sc-7208, 1:100), ERα (Abcam, Cat. #: ab37438, 1:200), Alexa Fluor (AF)488 K8 (Abcam, clone EP1628Y, Cat. #: ab192467, 1:200), K14 (Covance, Cat. #: PRB-155P, 1:400), TRANCE/TNFSF11/RANKL (R&D Biosystems, Cat. # AF462, 1:200), RANKL (Amgen, Cat. #: M366), K8+18 (Fitzgerald, Cat. #: 20R-CP004, 1:400), K5 (Fitzgerald, Cat. #: 20R-CP003, 1:400), RANK (Amgen, Cat. #: N1H8, 1:200), RANK (Santa Cruz Biotechnology, clone H-7, Cat. #: sc-374360, 1:100), and RANK (R&D Biosystems, Cat. # AF692, 1:200). Briefly, frozen sections were fixed in 2% paraformaldehyde, prior to permeabilization with 3% triton-x-100 and incubation with primary antibodies overnight at 4 °C. The next morning, sections were incubated with species specific secondary antibodies conjugated to different fluorophores (AF-633, -555, -568, -488, Invitrogen). All washes were carried out using Tris-buffered saline with 0.5% Tween-20 and nuclei and/or actin filaments were counterstained using 4′,6-diamidino-2-phenylindole (DAPI, Cat. #: D1306) or the appropriate Phalloidin-fluorophore conjugate (AF488, AF555, AF647, Cat. #’s: A12379, A34055 and A22287), respectively. Images of stained sections were acquired either on: an inverted Eclipse Ti-E Nikon microscope with CSU-X1 spinning disk confocal (Yokogawa Electric Corporation), 405 nm, 488 nm, 561, 635 nm lasers; a Plan Apo VC 60X/1.40 Oil or an Apo LWD 40X/1.15 Water-immersion λS objective; electronic shutters; a charge-coupled device (CCD) camera (Clara; Andor) and controlled by Metamorph; or a Nikon Ti Microscope (Inverted) with CSU-W1 widefield spinning disk confocal (Andor Borealis), 100 mW at 405, 561, and 640 nm; 150 mW at 488 nm lasers, an Andor Zyla sCMOS camera (5.5 megapixels) and Andor iXon Ultra DU888 1k x 1k Electron multiplying CCD to enable large field of view confocal imaging; and controlled by Micromanager.

### Image Analysis

Image analysis of IHC and immunofluorescence was performed using ImageJ and QuPath software^114,115^. For comparison, immunofluorescence images were subjected to same-level thresholding based on a determined range of positive fluorescence intensity in each channel and antibody staining panel and the threshold area (μm) was expressed as a percentage of whole cell or nuclear area using Phalloidin or DAPI staining measured in the same manner. Other IHC analysis was performed using the IHC profiler ImageJ plugin as described^116^.

### Fluorescence activated cell sorting (FACS) and primary MEC isolation

Tissues were digested as described abovem spun down at 1200 rpm and washed with FACS wash buffer (PBS with 2% FBS). Cells were then blocked with mouse and rat serum (Jackson Immunoresearch Laboratories, Cat. #: AB_2337194 and AB_2337141) and TruStain FcX (anti-mouse CD16/32) Antibody (BioLegend, clone 93, Cat. #: 101319) for 10 min, followed by incubation for 25 min at 4° C with the following mouse specific antibodies to delineate subpopulations: CD24-PE (BD Biosciences, clone M1/69, Cat, #: 553262, 1:100), CD49f -PE-Cy7 (BioLegend, clone GoH3, Cat: #: 313621, 1:100), CD31-APC (BioLegend, clone 390, Cat: #: 102409, 1:40), CD45-APC (BioLegend, clone 30-F11, Cat: #: 103111, 1:160), and TER-119-APC (BioLegend, clone TER-119, Cat: #: 116211, 1:80) and CD61-FITC (BioLegend, clone 2C9.G2 (HMβ3-1), Cat. #: 104305, 1:20). Human tissues were incubated for 25 min at 4 °C with the following primary antibodies: CD24-PE (BioLegend, clone ML5, Cat, #: 311105, 1:100), CD44-FITC (BioLegend, clone IM7, Cat: #: 103021, 1:20), EPCAM CD326-FITC (Invitrogen, clone VU-1D9, Cat. #: A15755, 1:100), CD49f-PE-Cy7 (BioLegend, clone GoH3, Cat: #: 313621, 1:100), CD31-PE (BioLegend, clone WM59, Cat: #: 303105, 1:40), CD45-PE (BioLegend, clone H130, Cat: #: 304007, 1:100) and CD235-α-PE (BD Biosciences, clone GA-R2, Cat: #: 561051, 1:100). MEC lineage negative populations were removed by positive APC-CD45/CD31/TER-119 (mouse) and PE-CD45/CD31/CD235-α (human) staining. Cells were washed with FACS wash buffer and with DAPI in PBS for 5 min to distinguish live/dead cells. BD FACSAria II cell sorters were used to conduct cell sorting using FACSDiva software (BD Biosceinces). Data was analyzed using FlowJo software (Tree Star). Isolated cells were collected and used for RNA isolation, colony formation assays or limiting dilution transplantation assays as described above.

### Mammary gland wholemounts

Inguinal mammary glands wholemounts were allowed to dry for 30 min prior to fixation and staining. For H&E staining, mammary glands were fixed in 4% paraformaldehyde prior to dehydration in xylene/alcohol series, H&E counterstain, and mounting with permount. For Carmine alum staining, glands were fixed with Carnoy’s solution (60% ethanol, 30% chloroform and 10% glacial acetic acid) and stained overnight in Carmine Alum (0.2% carmine and 0.5% potassium aluminum sulfate in water) prior to dehydration, clearing in xylene and mounting with permount.

### Limiting dilution transplantation assays (LDTAs)

LDTAs were performed as described^117^ using FACS isolated MECs. Briefly, CD24+CD49f^hi^ MECs (basal/myoepithelial) or PyMT tumor cells were counted such that increasing dilutions could be prepared (1000, 100, 50, 10 cells) for injection into a cleared inguinal mammary fat pad. The inguinal mammary fat pads of 3–4-week-old mice were cleared of their endogenous epithelium by removing the portion of the fat pad proximal to the lymph node. For normal MECs, the basal population was double sorted prior to cell transplantation. Clearance was verified by mounting resected glands on histology slides, fixing in Carnoy’s solution and staining with Carmine alum. Stem cell frequency was determined at 6 weeks post transplantation by the appearance of progressing tumors (PyMT tumor cells) or by examining mammary glands for epithelial ductal outgrowths as determined upon their harvest for mounting, fixation and carmine aluminum staining as described. Repopulating events were scored as positive outgrowths with evidence of ductal branching. For PyMT tumor LDTAs, three tumors each of the primary LDTAs taken from SOFT and STIFF ECM stroma conditions were pooled before counting cells and repeating the LDTA for secondary tumor formation. Tumor initiating cell and mammary epithelial progenitor cell frequency were determined by counting the number of positive outgrowths (tumor or ductal epithelium) from transplants made at each cell dilution and using an Extreme Limiting Dilution Analysis software application webtool from The Walter and Eliza Hall Institute of Medical Research (http://bioinf.wehi.edu.au/software/elda/)^118^.

### Atomic Force Microscopy (AFM)

AFM was performed and analyzed as described^17^ using an MFP3D-BIO inverted optical AFM (Asylum Research, Santa Barbara, CA) mounted on a Nikon TE200-U inverted microscope (Melville, NY) and placed on a vibration-isolation table (Herzan TS-150). Briefly, tissue specimens were allowed to thaw and equilibrate to room temperature before immersion in PBS supplemented with 1 ug/mL propidium iodide, protease and phosphatase inhibitors. A 2-μm beaded tip attached to a silicon nitride cantilever (Asylum Research, Santa Barbara, CA) was used for indentation. The spring constant of the cantilever was ∼ 0.06 N/m and cantilevers were calibrated using the thermal fluctuation method. AFM force maps were performed on 40 x 40 μm fields from two different quadrants of the same non-malignant human breast specimen, or as single points of indentation in a grid pattern for two regions from a HER2-positive breast cancer patient specimen. Elastic moduli measurements were derived from force curves obtained utilizing the FMAP function of the Igor Pro v. 6.22A (WaveMetrics, Lake Oswego, OR) supplied by Asylum Research. Cells were assumed to be incompressible and a Poisson’s ratio of 0.5 was used in the calculation of the Young’s elastic modulus. AFM measurements were generated for up to one hour after thawing tissues. Periductal stromal ECM-rich regions were selected to generate all force maps and single point indentations for each patient specimen.

### Second Harmonic Generation Imaging / 2-photon Microscopy

2-photon microscopy and second harmonic generation (SHG) imaging of collagen fiber thickness and orientation was performed as previously described^17,18^.

### Vaginal cytology for estrus cycle determination

The estrus cycle in mice was determined by vaginal cytology as described^62^. Briefly, a thin cotton tipped applicator (Puritan, Cat. #: 25-826 5WC) was immersed in sterile PBS prior to insertion into the vaginal cavity to collect fluid. The cotton tipped applicator was then smeared onto the well of a 24-well tissue culture plate before staining cells with crystal violet solution (0.2%). Wells were then examined under brightfield with an Olympus microscope (IX81) to determine the stage of estrus (proestrus, diestrus, metestrus and estrus) for each mouse by the cell content and morphology present.

### Primary cell colony formation assays

Primary FACS-isolated mouse luminal and basal MECs were plated at a density of 5000 cells and 1000 cells respectively in ∼20uL of 100% rBM (R&D Systems, Cultrex BME) by pipetting around the edge of the well of a 96 well plate. Cell Medium (DMEM-F12) containing 1% FBS, 500 ng/ml hydrocortisone, 5 μg/ml insulin and 20 ng/ml EGF was added to each well and colonies were counted after 10-14 days in culture with media exchanges every 2-3 days. In some cases, primary colonies were harvested from rBM gels using cell recovery solution (Corning, Cat. #: 354253), dissociated to single cells with Trypsin/EDTA, counted and then plated at the above densities to assess their proficiency at secondary colony formation. Resulting colonies were again counted after 10-14 days of culture with media exchanges every 2-3 days. For human tissues, fresh FACS isolated luminal and basal MECs were resuspended in rBM at a density of at 50,000 cells/mL, and 20-μl droplets (1000 cells) were seeded into 8-well chamber slides (LabTek II, Cat. #: 154534) and allowed to set at 37 °C. Wells were filled 0.4 mL cell medium (DMEM/F-12) containing Glutamax, B27 Supplement (Gibco, Cat. #: 17504044), 500 ng/ml hydrocortisone, 5 μg/ml insulin, 10 ng/ml EGF and 20 ng/ml Cholera Toxin from *Vibrio cholerae* (Sigma, C8052). Cells were cultured for 7-10 days prior to counting the number of colonies per well.

### Enzyme-linked immunosorbent assay (ELISA)

ELISAs were performed using Mouse TRANCE/RANKL/TNFSF11 DuoSet ELISA Kits (R&D Biosystems, Cat. #: DY462) according to the manufacturer’s directions.

### Mammography

All mammography was conducted by licensed physicians at the UCSF and Duke medical centers according to established protocols as described^48,119^. Quantitative measurements of the raw mammography images used the automated volumetric density measure developed by Dr. John Shepherd^120^.

### Breast cancer Patient Derived Xenografts (PDXs)

PDX tissues were obtained from Dr. Welm at the Huntsman Cancer Institute, University of Utah, Utah (HCI-012) or Dr. Michael Lewis at Baylor College of Medicine, San Antonio, Texas (BCM-3143B and BCM-3963)^57,58^. PDX fragments were established from frozen and maintained in NOD-SCID immunodeficient mice. Tumor initiation was a measure of how efficiently PDX fragments established as progressing tumors in mice upon primary implantation from cryopreserved tumor tissue. Once established tumors reached experimental endpoint, mice were sacrificed, and tumor tissue fragments were orthotopically transplanted into NOD-SCID recipient mice. The remaining tumor tissue was divided into pieces for formalin fixation and paraffin embedding, embedding and freezing in OCT, flash freezing and cryopreservation in 95% FBS: 5% DMSO. Flash frozen tumor pieces were used for RNA and protein isolation for the downstream applications indicated.

### Cell fractionation, Immunoblotting and Densitometry

When indicated, cell lysates were prepared to fractionate nuclear and cytoplasmic cellular compartments using Thermo Scientific NE-PER Nuclear and Cytoplasmic Extraction Reagents (Cat. #: 78833). Other cell lysates were prepared using RIPA buffer (150 mM NaCl, 1% NP40, 0.5% sodium deoxycholate, 0.1% sodium dodecyl sulphate, 50 mM Tris-HCl, pH 8.0) and all lysates were supplemented with protease and phosphatase inhibitor cocktail solutions (genDEPOT, Xpert solutions, Cat. #: P3100 and P3200). Immunoblotting was performed as described^121^ using Immobilon P Polyvinylidene difluoride membrane (PVDF, Millipore, Cat. #: IPVH00010) for protein transfer, and incubation with the following primary antibodies: phospho-p44/42 MAPK (EER1/2) (T202/Y204) (Cell Signaling Technology, Cat. #: 9101, 1:1000), p44/42 MAPK (ERK1/2) (Cell Signaling Technology, Cat. #: 9102, 1:1000), phospho-PR (S294) (Invitrogen, Cat. #: PA5-37472, 1:200), phospho-PR (S294) (Invitrogen, Cat. #: MA1-414, 1:200), PR (Santa Cruz Biotechnology, clone H-190, Cat. #: sc-7208, 1:500), NFκB-p65 (D14E12) XP (Cell Signaling Technology, Cat. #: 8242, 1:1000), and Lamin B1 (Abcam, Cat. #: ab16048, 1:2000). Immunoblots were then incubated with the appropriate species specific HRP-conjugated secondary antibodies (Advansta) before developing reactivity with pierce enhanced chemiluminescence (ECL) reagent (Advansta WesternBright ECL HRP substrate, Cat. #: K-12045). Membrane signals were visualized with a PXi imaging system (Syngene). Densitometry measurements were made using ImageJ software with normalization to background membrane density, the corresponding total protein levels, as well as levels of the loading controls LaminB1 and GAPDH.

### Cell Culture and treatments

T47D and MCF7 human breast cancer cell lines were obtained from the American Type Culture Collection (ATCC) and cultured as recommended in RPMI with 10% FBS and DMEM with 10% FBS, respectively. Cells were cultured on polyacrylamide gels of low and high stiffnesses (400 Pa and 6 kPa), serum starved overnight and then treated for 15 min with R5020 (10 nM) and EGF (20 ng/mL) alone or in combination. For inhibitor studies, the MEK inhibitor (U0126; 12.5μM, LCLabs, Cat. #: NC0976210) was incubated with cells for 1hr prior to R5020 and EGF stimulation.

### Polyacrylamide hydrogels

Polyacrylamide hydrogels of varying rigidities were prepared as described^122,123^. Briefly, compliant Polyacrylamide hydrogels were polymerized on silanized coverslips and functionalized with Fibronectin (1μg/ml) overnight. PA gels were washed three times and equilibrated overnight with cell media at 37 °C prior to cell seeding. Cells were cultured for 24 hr on hydrogels prior to treatment and cell lysis for immunoblotting.

### Statistical analysis

GraphPad Prism Version 9.1.2 was used to perform all statistical analyses and statistical significance was determined using the appropriate tests as noted in the corresponding figure legends. Tests of normality were used to determine the appropriate statistical test. All independent variables are described in the text with measurements always from distinct samples (biological replicates) unless otherwise stated. All tests are two-tailed unless otherwise indicated.

## DATA AVAILABILITY

The authors declare that all data supporting the findings of this study are available within this publication and its extended data. The RNAseq data has been deposited in NCBI’s Gene Expression Omnibus^124^ and are accessible through GEO Series accession number GSE179983 (https://www.ncbi.nlm.nih.gov/geo/query/acc.cgi?acc=GSE179983).

## CODE AVAILABILITY STATEMENT

All code used in the preparation of this manuscript is publicly available from software and commercial sources.

## ACKNOWLEDGEMENTS

We thank Lydia and Nataliya Korets for care and handling of animals. We also thank Nataliya Korets for tissue histology, the Nikon Imaging Center and Biological Imaging Development Center for microscopy support, as well as the Parnassus Flow Cytometry Core for support with cell sorting. We also thank Susan Samson for editorial input. This work was supported by US National Institutes of Health/National Cancer Institute grants 1R01CA222508-01 (V.M.W., E.S.H.), R01CA192914 and 1R35CA242447-01A1 (V.M.W), US Department of Defense (DOD) Breast Cancer Research Program (BCRP) grant BC122990 (V.M.W.), Give Breast Cancer the Boot pilot project grant (V.M.W, R.M.), and American Association for Cancer Research grants 15-40-01-NORT: Basic Cancer Research Fellowship (J.J.N.), and 17-40-48-NORT: AACR Janssen Cancer Interception Research Fellowship (J.J.N.).

## CONTRIBUTIONS

V.M.W. conceived and designed the study. V.M.W and J.J.N directed the studies. J.J.N., F.K. and C.S. conducted AFM analysis. C.S. performed RNAseq and gene set enrichment analyses. J.J.N. and M-K.H. performed H&E and IHC staining and qRT-PCR analysis of mouse and human breast tissues. A.J.I. assessed mouse and human tissue pathology. J.J.N. and D.T. performed T47D and MCF7 polyacrylamide hydrogel culture, hormone treatment and immunoblotting. J.J.N and Y.Y. completed all animal studies. J.J.N. performed FACS isolation of MECs, lineage tracing, colony formation and limiting dilution transplantation assays. J.J.N. performed collagen/rBM hydrogel studies *in vitro* and *in vivo* with PyMT tumor cells and PDX tissues. J.K.M participated in study design and data analysis. J.N.L. designed and developed the V737N mutant mouse model and provided technical support. S.S. provided intellectual input on study design and editorial input. R.A.M. and E.S.H. collected and provided human breast tissue specimens with patient data and contributed intellectually to study design and analysis. V.M.W. and J.J.N. wrote the manuscript with editorial input from all authors.

## REFERENCES

1. Tomasetti, C., Li, L. & Vogelstein, B. Stem cell divisions, somatic mutations, cancer etiology, and cancer prevention. Science 355, 1330–1334 (2017).

2. Tomasetti, C. & Vogelstein, B. Cancer etiology. Variation in cancer risk among tissues can be explained by the number of stem cell divisions. Science 347, 78–81 (2015).

3. Blokzijl, F., et al. Tissue-specific mutation accumulation in human adult stem cells during life. Nature 538, 260–264 (2016).

4. Shibue, T. & Weinberg, R.A.EMT, CSCs, and drug resistance: the mechanistic link and clinical implications. Nat Rev Clin Oncol 14, 611–629 (2017).

5. Taube, J.H., et al. Core epithelial-to-mesenchymal transition interactome gene-expression signature is associated with claudin-low and metaplastic breast cancer subtypes. Proc Natl Acad Sci U S A 107, 15449–15454 (2010).

6. Prat, A., et al. Phenotypic and molecular characterization of the claudin-low intrinsic subtype of breast cancer. Breast Cancer Res 12, R68 (2010).

7. Bierie, B., et al. Integrin-beta4 identifies cancer stem cell-enriched populations of partially mesenchymal carcinoma cells. Proc Natl Acad Sci U S A 114, E2337–E2346 (2017).

8. Glinsky, G.V. “Stemness” genomics law governs clinical behavior of human cancer: implications for decision making in disease management. J Clin Oncol 26, 2846–2853 (2008).

9. Northey, J.J., Przybyla, L. & Weaver, V.M. Tissue Force Programs Cell Fate and Tumor Aggression. Cancer Discov 7, 1224–1237 (2017).

10. Visvader, J.E. Cells of origin in cancer. Nature 469, 314–322 (2011).

11. Hayward, M.K., Muncie, J.M. & Weaver, V.M. Tissue mechanics in stem cell fate, development, and cancer. Dev Cell (2021).

12. Engler, A.J., Sen, S., Sweeney, H.L. & Discher, D.E. Matrix elasticity directs stem cell lineage specification. Cell 126, 677–689 (2006).

13. Engler, A.J., et al. Myotubes differentiate optimally on substrates with tissue-like stiffness: pathological implications for soft or stiff microenvironments. The Journal of cell biology 166, 877–887 (2004).

14. Przybyla, L., Lakins, J.N. & Weaver, V.M. Tissue Mechanics Orchestrate Wnt-Dependent Human Embryonic Stem Cell Differentiation. Cell Stem Cell 19, 462–475 (2016).

15. Muncie, J.M., et al. Mechanical Tension Promotes Formation of Gastrulation-like Nodes and Patterns Mesoderm Specification in Human Embryonic Stem Cells. Dev Cell 55, 679–694 e611 (2020).

16. Maller, O., et al. Tumour-associated macrophages drive stromal cell-dependent collagen crosslinking and stiffening to promote breast cancer aggression. Nat Mater 20, 548–559 (2021).

17. Acerbi, I., et al. Human breast cancer invasion and aggression correlates with ECM stiffening and immune cell infiltration. Integr Biol (Camb) 7, 1120–1134 (2015).

18. Pickup, M.W., et al. Stromally derived lysyl oxidase promotes metastasis of transforming growth factor-beta-deficient mouse mammary carcinomas. Cancer Res 73, 5336–5346 (2013).

19. Laklai, H., et al. Genotype tunes pancreatic ductal adenocarcinoma tissue tension to induce matricellular fibrosis and tumor progression. Nat Med 22, 497–505 (2016).

20. Guo, W., et al. Slug and Sox9 cooperatively determine the mammary stem cell state. Cell 148, 1015–1028 (2012).

21. Lambert, A.W. & Weinberg, R.A. Linking EMT programmes to normal and neoplastic epithelial stem cells. Nat Rev Cancer 21, 325–338 (2021).

22. Wilson, M.M., Weinberg, R.A., Lees, J.A. & Guen, V.J. Emerging Mechanisms by which EMT Programs Control Stemness. Trends Cancer 6, 775–780 (2020).

23. Lopez, J.I., Mouw, J.K. & Weaver, V.M. Biomechanical regulation of cell orientation and fate. Oncogene 27, 6981–6993 (2008).

24. Paszek, M.J., et al. Tensional homeostasis and the malignant phenotype. Cancer cell 8, 241–254 (2005).

25. Levental, K.R., et al. Matrix crosslinking forces tumor progression by enhancing integrin signaling. Cell 139, 891–906 (2009).

26. Mouw, J.K., et al. Tissue mechanics modulate microRNA-dependent PTEN expression to regulate malignant progression. Nat Med 20, 360–367 (2014).

27. Miroshnikova, Y.A., et al. Tissue mechanics promote IDH1-dependent HIF1alpha-tenascin C feedback to regulate glioblastoma aggression. Nat Cell Biol 18, 1336–1345 (2016).

28. Rubashkin, M.G., et al. Force engages vinculin and promotes tumor progression by enhancing PI3K activation of phosphatidylinositol (3,4,5)-triphosphate. Cancer Res 74, 4597–4611 (2014).

29. Desai, S.S., et al. Physiological ranges of matrix rigidity modulate primary mouse hepatocyte function in part through hepatocyte nuclear factor 4 alpha. Hepatology 64, 261–275 (2016).

30. Han, S., Pang, M.F. & Nelson, C.M. Substratum stiffness tunes proliferation downstream of Wnt3a in part by regulating integrin-linked kinase and frizzled-1. J Cell Sci 131(2018).

31. Storch, U., Mederos y Schnitzler, M. & Gudermann, T. G protein-mediated stretch reception. Am J Physiol Heart Circ Physiol 302, H1241–1249 (2012).

32. Samuel, M.S., et al. Actomyosin-mediated cellular tension drives increased tissue stiffness and beta-catenin activation to induce epidermal hyperplasia and tumor growth. Cancer Cell 19, 776–791 (2011).

33. Joshi, P.A., et al. Progesterone induces adult mammary stem cell expansion. Nature 465, 803–807 (2010).

34. Asselin-Labat, M.L., et al. Control of mammary stem cell function by steroid hormone signalling. Nature 465, 798–802 (2010).

35. Tanos, T., et al. Progesterone/RANKL is a major regulatory axis in the human breast. Sci Transl Med 5, 182ra155 (2013).

36. Yaghjyan, L., Colditz, G.A., Rosner, B. & Tamimi, R.M. Mammographic breast density and breast cancer risk by menopausal status, postmenopausal hormone use and a family history of breast cancer. Cancer Causes Control 23, 785–790 (2012).

37. Harvey, J.A., et al. Histologic changes in the breast with menopausal hormone therapy use: correlation with breast density, estrogen receptor, progesterone receptor, and proliferation indices. Menopause 15, 67–73 (2008).

38. Wood, C.E., et al. Progestin effects on cell proliferation pathways in the postmenopausal mammary gland. Breast Cancer Res 15, R62 (2013).

39. Boyd, N.F., et al. Mammographic density as a surrogate marker for the effects of hormone therapy on risk of breast cancer. Cancer Epidemiol Biomarkers Prev 15, 961–966 (2006).

40. Byrne, C., et al. Mammographic Density Change With Estrogen and Progestin Therapy and Breast Cancer Risk. J Natl Cancer Inst 109(2017).

41. Mulac-Jericevic, B., Lydon, J.P., DeMayo, F.J. & Conneely, O.M. Defective mammary gland morphogenesis in mice lacking the progesterone receptor B isoform. Proc Natl Acad Sci U S A 100, 9744–9749 (2003).

42. Aldaz, C.M., Liao, Q.Y., LaBate, M. & Johnston, D.A. Medroxyprogesterone acetate accelerates the development and increases the incidence of mouse mammary tumors induced by dimethylbenzanthracene. Carcinogenesis 17, 2069–2072 (1996).

43. Joshi, P.A., Goodwin, P.J. & Khokha, R. Progesterone Exposure and Breast Cancer Risk: Understanding the Biological Roots. JAMA oncology 1, 283–285 (2015).

44. Sampayo, R., Recouvreux, S. & Simian, M. The hyperplastic phenotype in PR-A and PR-B transgenic mice: lessons on the role of estrogen and progesterone receptors in the mouse mammary gland and breast cancer. Vitam Horm 93, 185–201 (2013).

45. Sprague, B.L., et al. Circulating serum xenoestrogens and mammographic breast density. Breast Cancer Res 15, R45 (2013).

46. Boyd, N.F., Martin, L.J., Yaffe, M.J. & Minkin, S. Mammographic density: a hormonally responsive risk factor for breast cancer. J Br Menopause Soc 12, 186–193 (2006).

47. Brisson, J., et al. Tamoxifen and mammographic breast densities. Cancer Epidemiol Biomarkers Prev 9, 911–915 (2000).

48. Northey, J.J., et al. Stiff stroma increases breast cancer risk by inducing the oncogene ZNF217. J Clin Invest 130, 5721–5737 (2020).

49. McConnell, J.C., et al. Increased peri-ductal collagen micro-organization may contribute to raised mammographic density. Breast Cancer Res 18, 5 (2016).

50. Winkler, J., Abisoye-Ogunniyan, A., Metcalf, K.J. & Werb, Z. Concepts of extracellular matrix remodelling in tumour progression and metastasis. Nat Commun 11, 5120 (2020).

51. Barnes, J.M., et al. A tension-mediated glycocalyx-integrin feedback loop promotes mesenchymal-like glioblastoma. Nat Cell Biol 20, 1203–1214 (2018).

52. Mekhdjian, A.H., et al. Integrin-mediated traction force enhances paxillin molecular associations and adhesion dynamics that increase the invasiveness of tumor cells into a three-dimensional extracellular matrix. Mol Biol Cell 28, 1467–1488 (2017).

53. Wei, S.C., et al. Matrix stiffness drives epithelial-mesenchymal transition and tumour metastasis through a TWIST1-G3BP2 mechanotransduction pathway. Nat Cell Biol 17, 678–688 (2015).

54. Hatzis, C., et al. A genomic predictor of response and survival following taxane-anthracycline chemotherapy for invasive breast cancer. JAMA 305, 1873–1881 (2011).

55. Girton, T.S., Oegema, T.R. & Tranquillo, R.T. Exploiting glycation to stiffen and strengthen tissue equivalents for tissue engineering. J Biomed Mater Res 46, 87–92 (1999).

56. Conklin, M.W., et al. Aligned collagen is a prognostic signature for survival in human breast carcinoma. Am J Pathol 178, 1221–1232 (2011).

57. Zhang, X., et al. A renewable tissue resource of phenotypically stable, biologically and ethnically diverse, patient-derived human breast cancer xenograft models. Cancer Res 73, 4885–4897 (2013).

58. DeRose, Y.S., et al. Patient-derived models of human breast cancer: protocols for in vitro and in vivo applications in tumor biology and translational medicine. Curr Protoc Pharmacol Chapter 14, Unit14 23 (2013).

59. Subramanian, A., et al. Gene set enrichment analysis: a knowledge-based approach for interpreting genome-wide expression profiles. Proc Natl Acad Sci U S A 102, 15545–15550 (2005).

60. Liberzon, A., et al. The Molecular Signatures Database (MSigDB) hallmark gene set collection. Cell Syst 1, 417–425 (2015).

61. van de Moosdijk, A.A., Fu, N.Y., Rios, A.C., Visvader, J.E. & van Amerongen, R. Lineage Tracing of Mammary Stem and Progenitor Cells. Methods Mol Biol 1501, 291–308 (2017).

62. Byers, S.L., Wiles, M.V., Dunn, S.L. & Taft, R.A. Mouse estrous cycle identification tool and images. PLoS One 7, e35538 (2012).

63. Rajaram, R.D., et al. Progesterone and Wnt4 control mammary stem cells via myoepithelial crosstalk. The EMBO journal 34, 641–652 (2015).

64. Dwyer, A.R., Truong, T.H., Ostrander, J.H. & Lange, C.A. 90 YEARS OF PROGESTERONE: Steroid receptors as MAPK signaling sensors in breast cancer: let the fates decide. J Mol Endocrinol 65, T35–T48 (2020).

65. Truong, T.H., et al. Phosphorylated Progesterone Receptor Isoforms Mediate Opposing Stem Cell and Proliferative Breast Cancer Cell Fates. Endocrinology 160, 430–446 (2019).

66. Knutson, T.P., et al. Phosphorylated and sumoylation-deficient progesterone receptors drive proliferative gene signatures during breast cancer progression. Breast Cancer Res 14, R95 (2012).

67. Knutson, T.P., et al. Posttranslationally modified progesterone receptors direct ligand-specific expression of breast cancer stem cell-associated gene programs. J Hematol Oncol 10, 89 (2017).

68. Lange, C.A., Shen, T. & Horwitz, K.B. Phosphorylation of human progesterone receptors at serine-294 by mitogen-activated protein kinase signals their degradation by the 26S proteasome. Proc Natl Acad Sci U S A 97, 1032–1037 (2000).

69. Duffy, S.W., et al. Mammographic density and breast cancer risk in breast screening assessment cases and women with a family history of breast cancer. Eur J Cancer 88, 48–56 (2018).

70. Lindstrom, S., et al. Genome-wide association study identifies multiple loci associated with both mammographic density and breast cancer risk. Nat Commun 5, 5303 (2014).

71. Boyd, N.F., et al. Mammographic density: a heritable risk factor for breast cancer. Methods Mol Biol 472, 343–360 (2009).

72. Gilbert, P.M., et al. Substrate elasticity regulates skeletal muscle stem cell self-renewal in culture. Science 329, 1078–1081 (2010).

73. Cosgrove, B.D., et al. Rejuvenation of the muscle stem cell population restores strength to injured aged muscles. Nat Med 20, 255–264 (2014).

74. Cozzolino, A.M., et al. Modulating the Substrate Stiffness to Manipulate Differentiation of Resident Liver Stem Cells and to Improve the Differentiation State of Hepatocytes. Stem Cells Int 2016, 5481493 (2016).

75. Lozoya, O.A., et al. Regulation of hepatic stem/progenitor phenotype by microenvironment stiffness in hydrogel models of the human liver stem cell niche. Biomaterials 32, 7389–7402 (2011).

76. Taddei, I., et al. Beta1 integrin deletion from the basal compartment of the mammary epithelium affects stem cells. Nat Cell Biol 10, 716–722 (2008).

77. Luo, M., et al. Distinct FAK activities determine progenitor and mammary stem cell characteristics. Cancer Res 73, 5591–5602 (2013).

78. Shibue, T., Brooks, M.W. & Weinberg, R.A. An integrin-linked machinery of cytoskeletal regulation that enables experimental tumor initiation and metastatic colonization. Cancer Cell 24, 481–498 (2013).

79. dos Santos, P.B., Zanetti, J.S., Ribeiro-Silva, A. & Beltrao, E.I. Beta 1 integrin predicts survival in breast cancer: a clinicopathological and immunohistochemical study. Diagn Pathol 7, 104 (2012).

80. Huck, L., Pontier, S.M., Zuo, D.M. & Muller, W.J. beta1-integrin is dispensable for the induction of ErbB2 mammary tumors but plays a critical role in the metastatic phase of tumor progression. Proc Natl Acad Sci U S A 107, 15559–15564 (2010).

81. Shibue, T. & Weinberg, R.A. Integrin beta1-focal adhesion kinase signaling directs the proliferation of metastatic cancer cells disseminated in the lungs. Proc Natl Acad Sci U S A 106, 10290–10295 (2009).

82. White, D.E., et al. Targeted disruption of beta1-integrin in a transgenic mouse model of human breast cancer reveals an essential role in mammary tumor induction. Cancer Cell 6, 159–170 (2004).

83. Timbrell, S., et al. FAK inhibition alone or in combination with adjuvant therapies reduces cancer stem cell activity. NPJ Breast Cancer 7, 65 (2021).

84. Lahlou, H., Sanguin-Gendreau, V., Frame, M.C. & Muller, W.J. Focal adhesion kinase contributes to proliferative potential of ErbB2 mammary tumour cells but is dispensable for ErbB2 mammary tumour induction in vivo. Breast Cancer Res 14, R36 (2012).

85. Provenzano, P.P. & Keely, P.J. The role of focal adhesion kinase in tumor initiation and progression. Cell Adh Migr 3, 347–350 (2009).

86. Provenzano, P.P., Inman, D.R., Eliceiri, K.W., Beggs, H.E. & Keely, P.J. Mammary epithelial-specific disruption of focal adhesion kinase retards tumor formation and metastasis in a transgenic mouse model of human breast cancer. Am J Pathol 173, 1551–1565 (2008).

87. Madan, R., Smolkin, M.B., Cocker, R., Fayyad, R. & Oktay, M.H. Focal adhesion proteins as markers of malignant transformation and prognostic indicators in breast carcinoma. Hum Pathol 37, 9–15 (2006).

88. Visvader, J.E. & Stingl, J. Mammary stem cells and the differentiation hierarchy: current status and perspectives. Genes Dev 28, 1143–1158 (2014).

89. Bhat, V., Lee-Wing, V., Hu, P. & Raouf, A. Isolation and characterization of a new basal-like luminal progenitor in human breast tissue. Stem Cell Res Ther 10, 269 (2019).

90. Callewaert, F., et al. Androgen receptor disruption increases the osteogenic response to mechanical loading in male mice. J Bone Miner Res 25, 124–131 (2010).

91. Gharahdaghi, N., et al. Links Between Testosterone, Oestrogen, and the Growth Hormone/Insulin-Like Growth Factor Axis and Resistance Exercise Muscle Adaptations. Front Physiol 11, 621226 (2020).

92. Page, A.J., et al. Ghrelin selectively reduces mechanosensitivity of upper gastrointestinal vagal afferents. Am J Physiol Gastrointest Liver Physiol 292, G1376–1384 (2007).

93. Kraemer, W.J. & Ratamess, N.A. Hormonal responses and adaptations to resistance exercise and training. Sports Med 35, 339–361 (2005).

94. Ataca, D., et al. The secreted protease Adamts18 links hormone action to activation of the mammary stem cell niche. Nat Commun 11, 1571 (2020).

95. Hosseini, H., et al. Early dissemination seeds metastasis in breast cancer. Nature 540, 552–558 (2016).

96. Mohammed, H., et al. Progesterone receptor modulates ERalpha action in breast cancer. Nature 523, 313–317 (2015).

97. Nolan, E., et al. RANK ligand as a potential target for breast cancer prevention in BRCA1-mutation carriers. Nat Med 22, 933–939 (2016).

98. Reyes, M.E., et al. Poor prognosis of patients with triple-negative breast cancer can be stratified by RANK and RANKL dual expression. Breast Cancer Res Treat 164, 57–67 (2017).

99. Pfitzner, B.M., et al. RANK expression as a prognostic and predictive marker in breast cancer. Breast Cancer Res Treat 145, 307–315 (2014).

100. Palafox, M., et al. RANK induces epithelial-mesenchymal transition and stemness in human mammary epithelial cells and promotes tumorigenesis and metastasis. Cancer Res 72, 2879–2888 (2012).

101. Gonzalez-Suarez, E., et al. RANK ligand mediates progestin-induced mammary epithelial proliferation and carcinogenesis. Nature 468, 103–107 (2010).

102. Huo, C.W., et al. High mammographic density is associated with an increase in stromal collagen and immune cells within the mammary epithelium. Breast Cancer Res 17, 79 (2015).

103. Sau, A., et al. Persistent Activation of NF-kappaB in BRCA1-Deficient Mammary Progenitors Drives Aberrant Proliferation and Accumulation of DNA Damage. Cell Stem Cell 19, 52–65 (2016).

104. Lim, E., et al. Aberrant luminal progenitors as the candidate target population for basal tumor development in BRCA1 mutation carriers. Nat Med 15, 907–913 (2009).

105. Moran, O., et al. Serum osteoprotegerin levels and mammographic density among high-risk women. Cancer Causes Control 29, 507–517 (2018).

106. Toriola, A.T., et al. Increased breast tissue receptor activator of nuclear factor-kappaB ligand (RANKL) gene expression is associated with higher mammographic density in premenopausal women. Oncotarget 8, 73787–73792 (2017).

107. Gonzalez-Suarez, E. & Sanz-Moreno, A. RANK as a therapeutic target in cancer. FEBS J 283, 2018–2033 (2016).

108. Tan, W., et al. Tumour-infiltrating regulatory T cells stimulate mammary cancer metastasis through RANKL-RANK signalling. Nature 470, 548–553 (2011).

109. Gomez-Aleza, C., et al. Inhibition of RANK signaling in breast cancer induces an anti-tumor immune response orchestrated by CD8+ T cells. Nat Commun 11, 6335 (2020).

110. Ahern, E., et al. RANKL blockade improves efficacy of PD1-PD-L1 blockade or dual PD1-PD-L1 and CTLA4 blockade in mouse models of cancer. Oncoimmunology 7, e1431088 (2018).

111. Yoldi, G., et al. RANK Signaling Blockade Reduces Breast Cancer Recurrence by Inducing Tumor Cell Differentiation. Cancer Res 76, 5857–5869 (2016).

112. Livet, J., et al. Transgenic strategies for combinatorial expression of fluorescent proteins in the nervous system. Nature 450, 56–62 (2007).

113. Hsu, H., et al. Tumor necrosis factor receptor family member RANK mediates osteoclast differentiation and activation induced by osteoprotegerin ligand. Proc Natl Acad Sci U S A 96, 3540–3545 (1999).

114. Bankhead, P., et al. QuPath: Open source software for digital pathology image analysis. Sci Rep 7, 16878 (2017).

115. Schindelin, J., et al. Fiji: an open-source platform for biological-image analysis. Nat Methods 9, 676–682 (2012).

116. Varghese, F., Bukhari, A.B., Malhotra, R. & De, A. IHC Profiler: an open source plugin for the quantitative evaluation and automated scoring of immunohistochemistry images of human tissue samples. PLoS One 9, e96801 (2014).

117. Shackleton, M., et al. Generation of a functional mammary gland from a single stem cell. Nature 439, 84–88 (2006).

118. Hu, Y. & Smyth, G.K. ELDA: extreme limiting dilution analysis for comparing depleted and enriched populations in stem cell and other assays. J Immunol Methods 347, 70–78 (2009).

119. Shepherd, J.A., et al. Clinical comparison of a novel breast DXA technique to mammographic density. Med Phys 33, 1490–1498 (2006).

120. Shepherd, J.A., et al. Volume of mammographic density and risk of breast cancer. Cancer Epidemiol Biomarkers Prev 20, 1473–1482 (2011).

121. Johnson, K.R., Leight, J.L. & Weaver, V.M. Demystifying the effects of a three-dimensional microenvironment in tissue morphogenesis. Methods Cell Biol 83, 547–583 (2007).

122. Przybyla, L., Lakins, J.N., Sunyer, R., Trepat, X. & Weaver, V.M. Monitoring developmental force distributions in reconstituted embryonic epithelia. Methods 94, 101–113 (2016).

123. Lakins, J.N., Chin, A.R. & Weaver, V.M. Exploring the link between human embryonic stem cell organization and fate using tension-calibrated extracellular matrix functionalized polyacrylamide gels. Methods Mol Biol 916, 317–350 (2012).

124. Edgar, R., Domrachev, M. & Lash, A.E. Gene Expression Omnibus: NCBI gene expression and hybridization array data repository. Nucleic Acids Res 30, 207–210 (2002).

