## Extended Data for "Mechanosensitive hormone signaling promotes mammary progenitor expansion and breast cancer progression"

EXTENDED DATA FIGURE LEGENDS

**Extended Data Fig. 1: V737N-β1 expression in the mammary gland promotes mechanosignaling and metastasis of Neu-induced tumors but has no effect on tumor latency or growth rate. a**, Schematic of strategy for V737N-β1−Integrin-IRES-EGFP transgene insertion at the Rosa26 locus of the mouse genome. **b**, (left) Representative FACS plot showing analysis of V737N-β1-Integrin expressing MECs using cell surface antigens, CD24 and CD49f, to examine GFP expression in the luminal (CD24+CD49f^lo^) and basal (CD24+CD49f^hi^) epithelial lineages. (right) Percentage of GFP-positive cells for each lineage are plotted (n=4 independent MEC isolations). **c**, Mammary glands harvested from wildtype (CTL) and V737N-β1-Integrin expressing mice (V737N) were fixed prior to embedding and freezing in OCT. Representative images of EGFP expression (green) in frozen tissue sections examined by confocal microscopy (n=3). Nuclei are stained with DAPI (cyan). Scale bar, 50 μm. **d**, Representative images of immunofluorescence staining for frozen tumor sections from Neu and V737N^Neu^ mice with an antibody specific to human β1-Integrin (red). Nuclei are stained with DAPI. Scale bar, 25 μm. **e**, (left) Representative images of immunofluorescence staining as in d (Neu; n=5, and V737N^Neu^; n=5) with an antibody specific to phospho-ERK (red; DAPI, cyan). Scale bar, 50 μm. (right) Average percentage of positive phospho-ERK staining per total cell area is plotted. **f**, (left) Representative images of immunofluorescence staining as in d (Neu; n=5, and V737N^Neu^; n=5) with an antibody specific to phosphorylated-AKT substrates (red; DAPI, cyan). Scale bar, 25 μm. (right) Average percentage of positive phospho-AKT substrates staining per total cell area is plotted. **g**, (left) Representative images of immunofluorescence staining as in d (Neu; n=5, and V737N^Neu^; n=5) with an antibody specific to phospho-FAK (red; DAPI, cyan). Scale bar, 50 μm. (right) Average percentage of positive phospho-FAK staining per total cell area is plotted. **h**, (left) Representative images of immunofluorescence staining and quantification as in d (Neu; n=5, and V737N^Neu^; n=5) with an antibody specific to active conformation β1-Integrin (magenta). Scale bar, 50 μm. Actin filaments were stained with Phalloidin (488, green). (right) Average percentage of positive active β1-integrin staining per total cell area is plotted. **i**, Graph of tumor latency (time to palpation) for Neu (n=13) and V737N^Neu^ (n=19) mice. **j**, Average tumor volume for Neu (n=13) and V737N^Neu^ (n=19) tumors as determined by weekly caliper measurement. **k**, (left) Representative images of immunofluorescence staining as in d (Neu; n=5, and V737N^Neu^; n=5) with an antibody specific to phospho-Histone H3 (red; DAPI, cyan). Scale bar, 25 μm. (right) Average percentage of phospho-Histone H3 positive cells are plotted. **l**, (left) Representative images of immunofluorescence staining as in d (Neu; n=5, and V737N^Neu^; n=5) with an antibody specific to cleaved Caspase-3 (red; DAPI, cyan). Scale bar, 25 μm. (right) Average percentage of cleaved Caspase-3 positive cells are plotted. **m**, Percentage of Neu (n=13) and V737N^Neu^ (n=19) mice with detectable lung metastases. **n-o**, Graphs showing qRT-PCR analysis of RNA isolated from Neu and V737N^Neu^ tumors displaying relative gene expression for the indicated genes. All graphs are presented as mean +/- S.E.M. Statistical tests used were Mann-Whitney test (b, f-h, k, l), unpaired *t*-test (e, n, o), log-rank test (i) and two-way ANOVA (with Bonferroni’s multiple comparisons test) (j). *P<0.03, **P<0.002, ***P<0.0002, ns=non-significant.

**Extended Data Fig. 2: A stiff extracellular matrix promotes mechanosignaling, metastasis and mesenchymal, tumor initiating cell frequency in PyMT tumors. a**, (left) Representative images of immunofluorescence staining for frozen sections from PyMT orthotopic tumors inoculated within SOFT (no L-ribose; n=5) and STIFF (L-ribose mediated crosslinking; n=5) Col1/rBM matrices using an antibody specific to active conformation β1-Integrin (magenta). Actin filaments are stained with Phalloidin (488, green). Scale bar, 50 μm. (right) Average percentage of positive active β1-Integrin staining per total cell area is plotted. **b**, Average tumor growth for PyMT orthotopic tumors in SOFT (n=8) and STIFF (n=8) matrices as determined by caliper measurement. **c**, Percentage of mice bearing PyMT orthotopic tumors in SOFT (n=8) and STIFF (n=8) matrices with detectable lung metastases. **d-e**, Graphs showing qRT-PCR analysis of RNA extracted from PyMT orthotopic tumors in SOFT (n=4) and STIFF (n=4) matrices showing relative gene expression for the indicated genes. **f**, Tumor initiating cell frequencies measured for SOFT and STIFF PyMT tumors (from a pool of three individual tumors each) by primary and secondary limiting dilution transplantation tumorigenesis assays in syngeneic mice. Results for primary assays are shown and correspond to secondary assays presented in main Figure 2. All graphs are presented as mean +/- S.E.M. Statistical tests used were Mann-Whitney test (a), unpaired *t*-test (d, e) and two-way ANOVA (with Bonferroni’s multiple comparisons test) (b). *P<0.03, **P<0.002, ***P<0.0002, ns=non-significant.

**Extended Data Fig. 3: A stiff stroma enhances mechanosignaling, tumor growth, metastasis and mesenchymal gene expression in HER2-positive breast cancer patient-derived xenografts. a**, (left) Representative images of immunofluorescence staining for frozen sections from HER2-positive breast cancer patient-derived xenografts (PDX) transplanted with SOFT (no L-ribose, n=6) and STIFF (L-ribose mediated crosslinking, n=6) Col1/rBM matrices using an antibody specific to active conformation β1-Integrin (magenta). Actin filaments are stained with Phalloidin (488, green). Scale bar, 50 μm. (right) Average percentage of positive active β1-Integrin staining per total cell area is plotted. **b-d**, Graphs showing average tumor growth in SOFT and STIFF matrices for the HER2-positive PDX models indicated as determined by caliper measurement (SOFT and STIFF, n=10 each for BCM-3963 and BCM3143B, n=4 each for HCI-012). **e-f**, Percentage of mice bearing HER2-positive PDX tumors with SOFT and STIFF matrices presenting detectable lung metastases (SOFT and STIFF, n=10 each for BCM-3963 and BCM3143B). **g-l**, Graphs showing qRT-PCR analysis of RNA extracted from HER2-positive PDX tumors with SOFT (n=7) and STIFF (n=7) matrices showing relative gene expression for the indicated mesenchymal and epithelial genes. All graphs are presented as mean +/- S.E.M. Statistical tests used were Mann-Whitney test (g-l), unpaired *t*-test (a) and two-way ANOVA (with Bonferroni’s multiple comparisons test) (b-d). *P<0.03, **P<0.002, ***P<0.0002, ns=non-significant.

**Extended Data Fig. 4: High mechanosignaling in the mouse mammary gland promotes precocious ductal branching and proliferation. a**, Representative images of immunofluorescence staining for frozen mammary gland sections from 10-week-old CTL and V737N mice with an antibody specific to human β1-Integrin (red). Nuclei are stained with DAPI (cyan). Scale bar, 25 μm. **b**, (left) Representative images of immunofluorescence staining as in d with an antibody specific to phospho-FAK (red). Nuclei are stained with DAPI (cyan). (right) The percentage of high positive phospho-FAK staining is plotted (n=5 for CTL and V737N mice). Scale bar, 25 μm. **c**, (left) Representative images of immunofluorescence staining as in d with an antibody specific to phospho-p130-CAS (red). Nuclei are stained with DAPI (cyan). (right) The percentage of high positive phospho-p130CAS staining is plotted (n=5 for CTL and V737N mice). Scale bar, 25 μm. **d-e**, (left) Images of whole mammary glands for CTL and V737N mice at 6- (top, a) and 10-weeks (bottom, b) of age (n=8-13 for CTL and V737N mice). Scale bar, 2 cm. (right) Average mammary gland weight (mg) for each corresponding age is plotted. **f**, Images from a and b were used to trace the outline of mammary glands to estimate and plot average surface area (mm^2^) and overall size (n=5 for CTL and V737N mice). **g**-**h**, (left) Brightfield images of H&E-stained mammary gland whole mounts for CTL and V737N mice at 6- (top, g) and 10-weeks (bottom, h) of age. (middle and right) Primary and secondary ductal branching was quantified for glands at each stage of development and plotted (n=3-5 for CTL and V737N mice). **i**-**j**, Representative images of immunofluorescence staining as in d for mice at 6- (far left) and 10-weeks (middle right) of age with an antibody specific to phosphorylated-Histone H3 (red) (n=5 for CTL and V737N mice). Nuclei are stained with DAPI (cyan). Scale bar, 25 μm. (middle left and far right) The percentage of positive phospho-Histone H3 nuclei per total nuclei is plotted to the right of corresponding representative images. All graphs are presented as mean +/- S.E.M. Statistical tests used were unpaired *t*-tests. *P<0.03, **P<0.002, ***P<0.0002, ns=non-significant.

**Extended Data Fig. 5: Luminal and Basal specific gene expression of mammary epithelial cells. a-e**, Graphs showing qRT-PCR analysis of sorted (FACS) MEC populations from CTL and V737N mice (n=3-7 independent isolations) to assess relative gene expression for the indicated luminal and basal specific genes. For all graphs error bars represent S.E.M. and statistical tests used were paired two-way ANOVA (with two-stage linear step-up procedure of Benjamini, Krieger and Yekutieli for multiple comparisons test) (a-e). *P<0.05, **P<0.005, ***P<0.0005, ns=non-significant.

**Extended Data Fig. 6: Colony formation and lineage tracing of mammary epithelial cells with activated mechanosignaling. a**, MECs sorted from CTL and V737N mice using surface antigens CD24 and CD49f to distinguish luminal (CD24^+^; CD49f^lo^) and basal (CD24^+^; CD49f^hi^) lineages were subjected to rBM colony formation assays to assess progenitor activity. The bar graph shows primary colony formation determined after 10 days in culture. **b**, A single dose of tamoxifen (1.5mg per mouse) was used to induce V737N-β1 and Confetti reporter expression at 3-4 weeks of age in K5-positive epithelial cells (K5/creERT2). Frequency of distinct clones per ductal region is plotted. **c**, Graph showing the relative percentage of single cell, two-cell, and multicellular clones at 2 weeks post tamoxifen induction in the ductal epithelium of mammary glands harvested from mice as in b. All graphs are presented as mean +/- S.E.M. Statistical tests used were two-way ANOVA (with Bonferroni’s multiple comparisons test) (a) and unpaired *t*-test (b). *P<0.03, **P<0.002, ***P<0.0002, ns=non-significant.

**Extended Data Fig. 7: Epithelial-to-mesenchymal transition and stem/progenitor related gene expression of mammary epithelial cells. a-i**, Graphs showing qRT-PCR analysis of RNA from FACs-isolated luminal and basal lineage MEC populations from CTL and V737N mice (n=5-6 independent isolations) to assess relative gene expression for the indicated EMT and stem/progenitor associated genes. All graphs are presented as mean +/- S.E.M. Statistical tests used were paired two-way ANOVA (with two-stage linear step-up procedure of Benjamini, Krieger and Yekutieli for multiple comparisons test) (a-e). *P<0.05, **P<0.005, ***P<0.0005, ns=non-significant.

**Extended Data Fig. 8: Hormone signaling related gene expression of mammary epithelial cells. a-d**, Graphs showing qRT-PCR analysis of sorted (FACS) luminal and basal lineage MEC populations from CTL and V737N mice (n=4-8 independent isolations) to assess relative gene expression for the indicated hormone signaling associated genes. All graphs are presented as mean +/- S.E.M. Statistical tests used were paired two-way ANOVA (with two-stage linear step-up procedure of Benjamini, Krieger and Yekutieli for multiple comparisons test) (a-e). *P<0.05, **P<0.005, ***P<0.0005, ns=non-significant.

**Extended Data Fig. 9: Extracellular matrix stiffness potentiates growth factor induced progesterone receptor phosphorylation in breast cancer cells. a**, MCF7 breast cancer cells were cultured on fibronectin conjugated polyacrylamide gels of varying stiffness (0.4 and 6kPa), serum starved overnight, and then treated with EGF (20ng/mL), R5020 (10nM), U0126 (MEK inhibitor), EGF and R5020 together (E+R), EGF and U0126 (MEK inhibitor, E+U), or U0126 for 1hr prior to EGF and R5020 (E+R+U) for 15 min. Cells were then lysed for immunoblotting with antibodies specific to phospho-PR (S294), PR-A and PR-B, p-ERK (T202/204) and ERK (n=3 biological replicates). **b**, Bar graph of densitometry measured for PR-B phosphorylation relative to total PR-B from three independent experiments conducted as in a (SOFT=0.4kPa; STIFF=6kPa). All treatment conditions were normalized to the untreated, serum starved condition. **c**, Bar graph of the data from b represented as fold change of STIFF/SOFT for each treatment condition. All graphs are presented as mean +/- S.E.M. Statistical tests used were two-way ANOVA (with two-stage linear step-up procedure of Benjamini, Krieger and Yekutieli for multiple comparisons test) (b, c). *P<0.05, **P<0.005, ***P<0.0005, ns=non-significant.

**Extended Data Fig. 10: A stiff extracellular matrix and high mechanosignaling promote RANK activity in mammary tumors. a-b**, Graphs showing qRT-PCR analysis of relative gene expression for *WNT4* and *TNFRSF11A* (*RANK)* in HER2-positive PDX tumors developed in SOFT (no L-ribose, n=8) and STIFF (L-ribose mediated crosslinking, n=8) stromal matrices. **c**, (left) Representative images of immunofluorescence staining for frozen sections of mammary tumors as in a with an antibody specific to RANK (red) and nuclei stained by DAPI (cyan) (n=6 for SOFT and STIFF). Scale bar, 50 μm. (right) Average percentage of positive staining per total cell area is plotted. **d**, (left) Representative images of immunofluorescence staining for frozen sections of mammary tumors as in a with an antibody specific to p65-NFκB (red). Actin filaments are stained by Phalloidin (488, green) (n=6 for SOFT and STIFF). Scale bar, 50 μm. (right) Average percentage of positive p65-NFκB staining per total nuclear area is plotted. **e**, Graph showing qRT-PCR analysis of relative expression for *Tnfsf11 (Rankl)* in PyMT mammary tumors inoculated within SOFT (n=5) and STIFF (n=5) stromal matrices as in a. **f**, (left) Representative images of immunofluorescence staining for frozen sections of mammary tumors inoculated as in e with an antibody specific to RANKL (red). Nuclei were stained with DAPI (cyan) (n=6 for SOFT and STIFF). Scale bar, 50 μm. (right) Average percentage of positive RANKL staining per total cell area is plotted. **g-h**, Graphs showing qRT-PCR analysis of relative gene expression for *Wnt4* and *Tnfrsf11a* (*Rank)* in SOFT (n=5) and STIFF (n=5) PyMT mammary tumors. **i**, (left) Representative images of immunofluorescence staining for frozen sections of PyMT mammary tumors as in e with an antibody specific to RANK (red). Nuclei were stained with DAPI (cyan) (n=6 for SOFT and STIFF). Scale bar, 50 μm. (right) Average percentage of positive RANK staining per total cell area is plotted. **j**, (left) Representative images of immunofluorescence staining for frozen sections of mammary tumors as in e with an antibody specific to p65-NFκB (red). Actin filaments are stained by Phalloidin (488, green (n=6 for SOFT and STIFF). Scale bar, 50 μm. (right) Average percentage of positive p65-NFκB staining per total nuclear area is plotted. **k**, Graph showing qRT-PCR analysis of relative expression for *Tnfsf11* (*Rankl)* in Neu (n=6) and V737N^Neu^ (n=6) mammary tumors. **l**, (left) Representative images of immunofluorescence staining for frozen sections of mammary tumors as in k with an antibody specific to RANKL (red). Nuclei were stained with DAPI (cyan) (n=6 for Neu and V737N^Neu^). Scale bar, 50 μm. (right) Average percentage of positive RANKL staining per total cell area is plotted. **m-n**, Graphs showing qRT-PCR analysis of relative gene expression for *Wnt4* and *Tnfrsf11a* (*Rank)* in Neu (n=6) and V737N^Neu^ (n=6) mammary tumors. **c**, (left) Representative images of immunofluorescence staining for frozen sections of mammary tumors as in k with an antibody specific to RANK (red). Nuclei were stained with DAPI (cyan) (n=6 for Neu and V737N^Neu^). Scale bar, 50 um. (right) Average percentage of positive RANK staining per total cell area is plotted. **d**, (left) Representative images of immunofluorescence staining for frozen sections of mammary tumors as in k with an antibody specific to p65-NFκB (red). Actin filaments are stained by Phalloidin (488, green) (n=6 for Neu and V737N^Neu^). Scale bar, 50 μm. (right) Average percentage of positive p65-NFκB staining per total nuclear area is plotted. All graphs are presented as mean +/- S.E.M. Statistical tests used were unpaired *t*-test (e, g, h, m n) and Mann-Whitney test (a-d, f, i-l, o, p). *P<0.03, **P<0.002, ***P<0.0002, ns=non-significant.

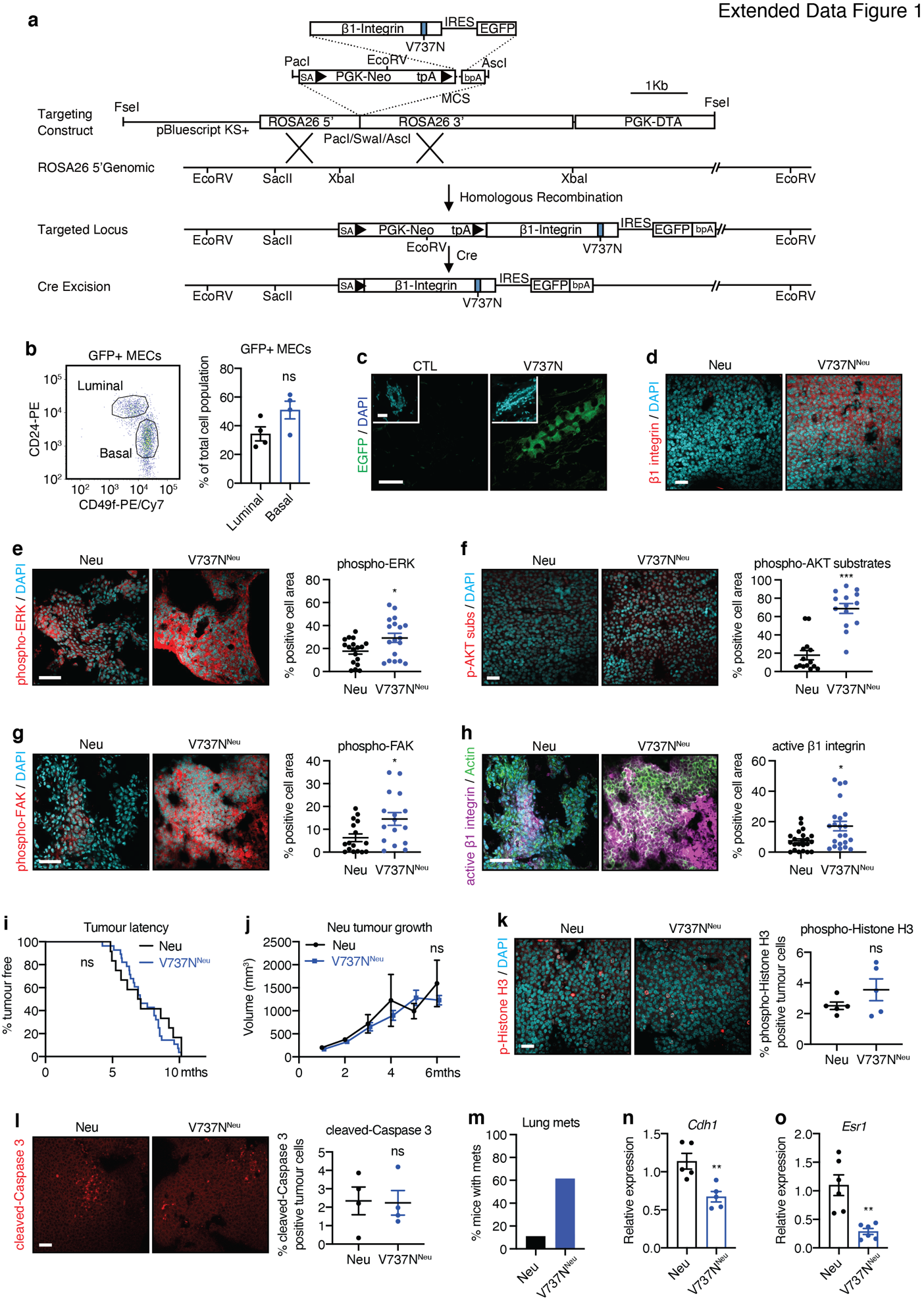

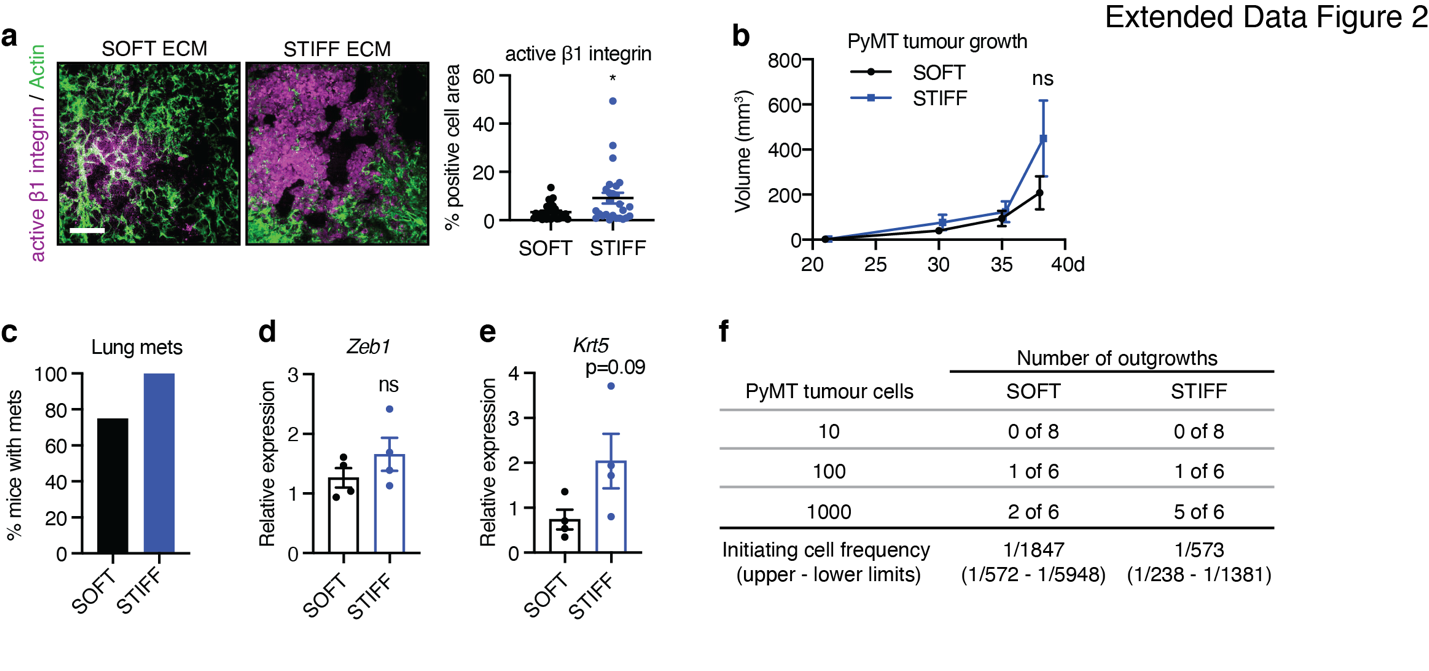

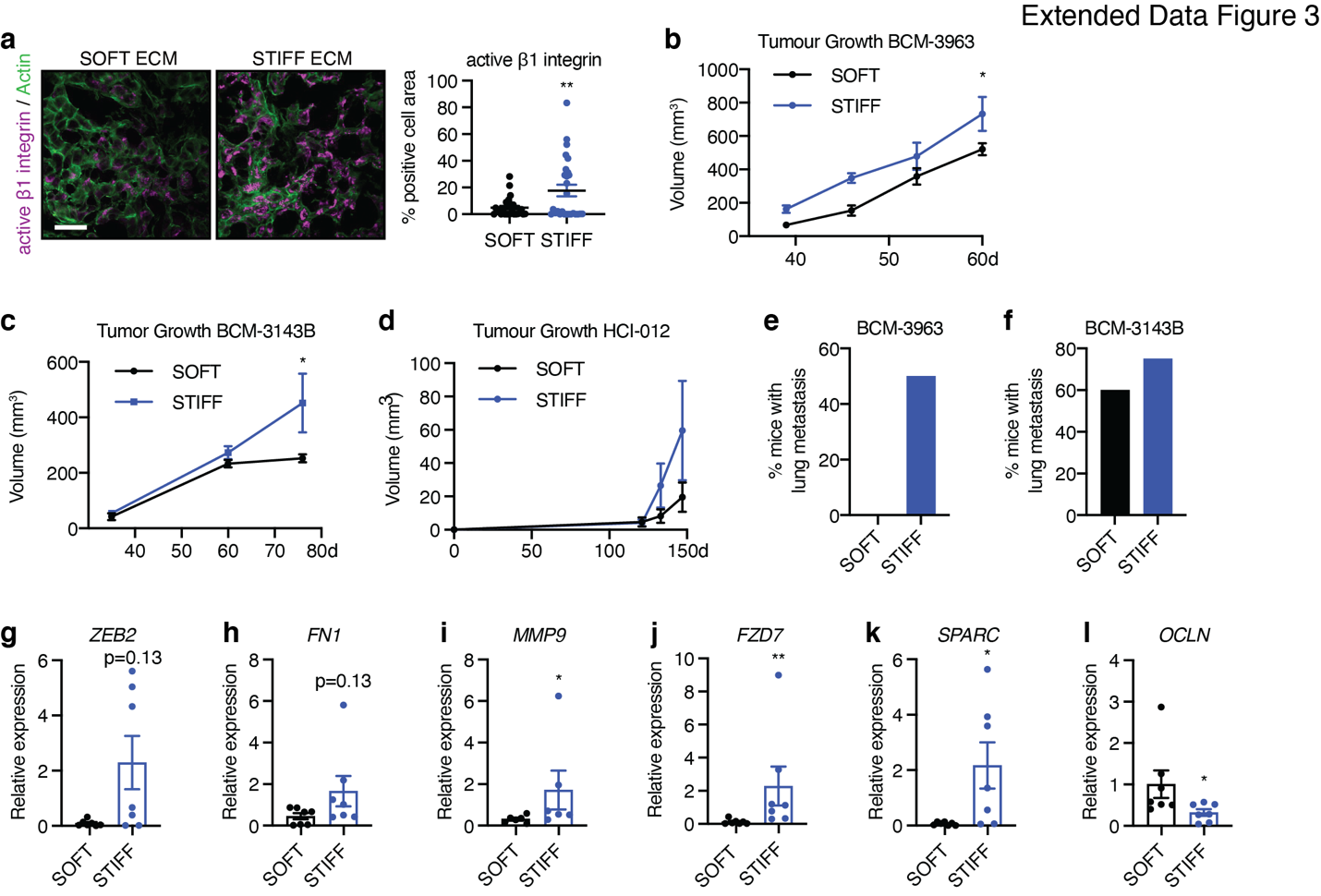

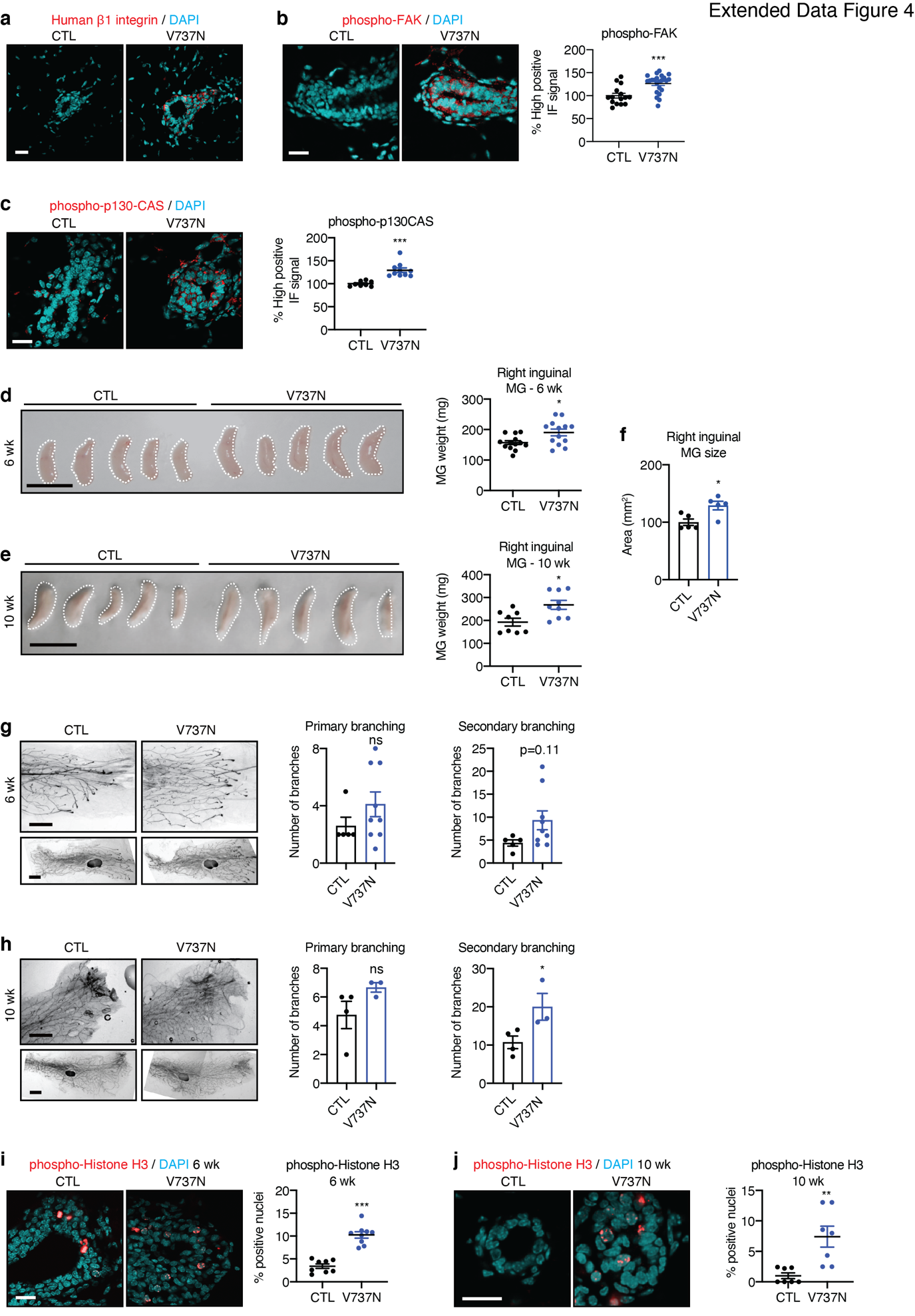

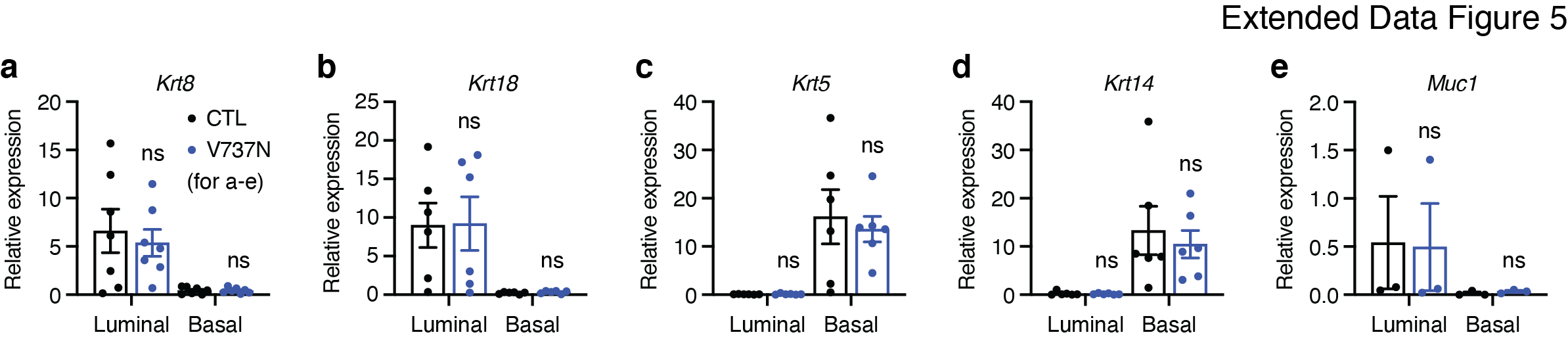

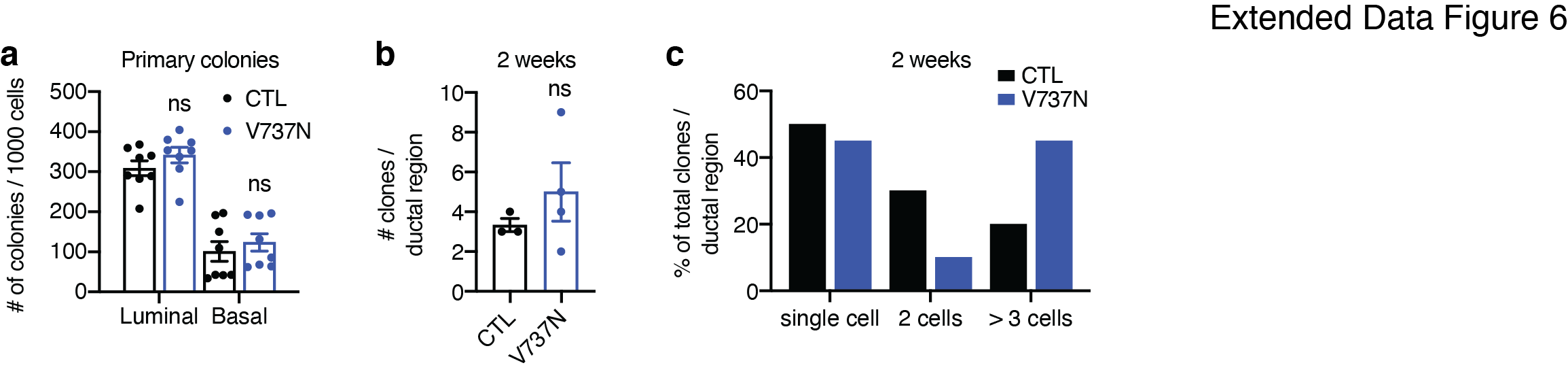

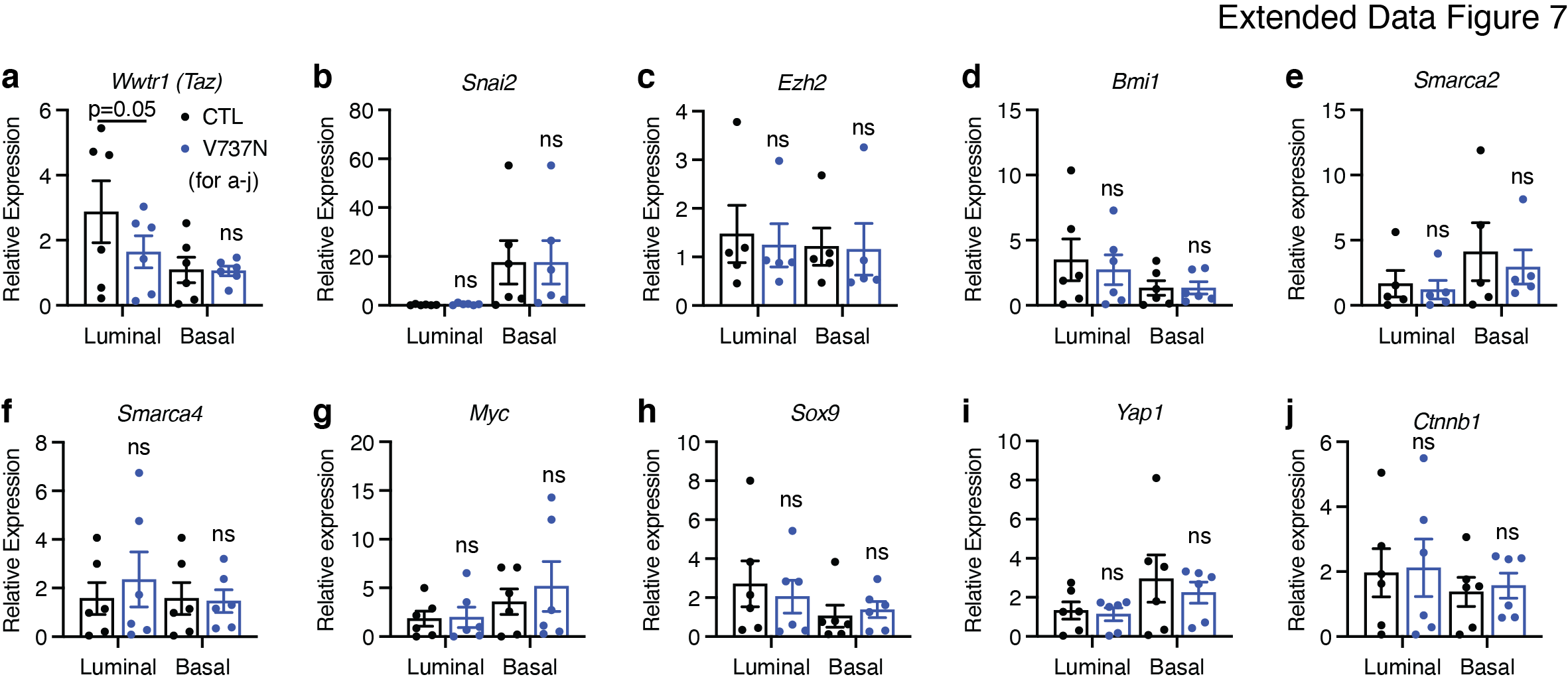

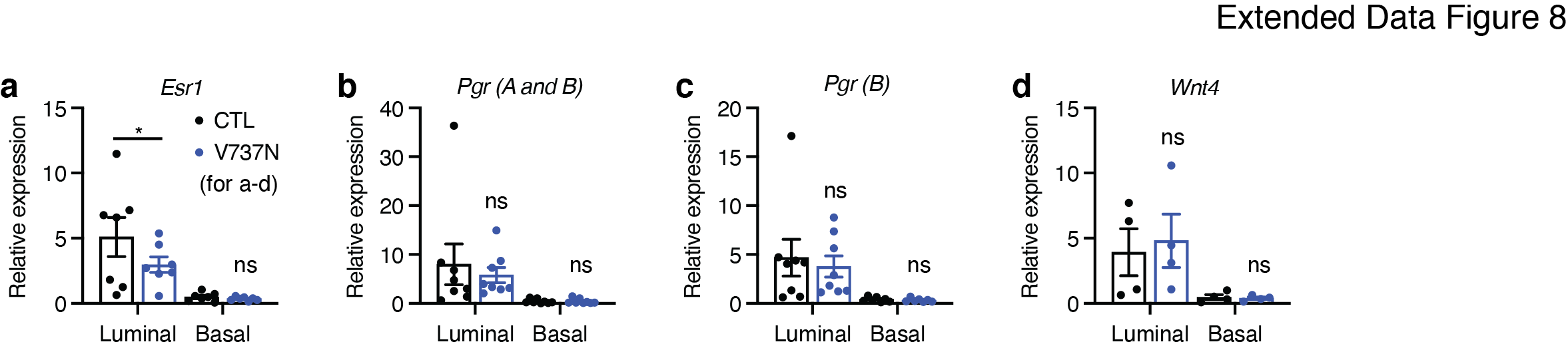

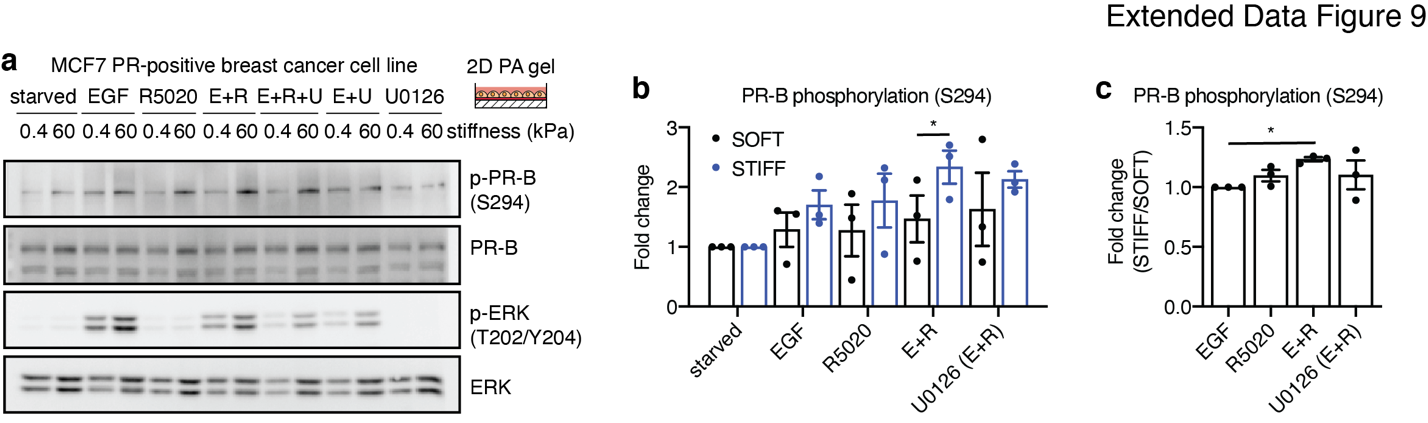

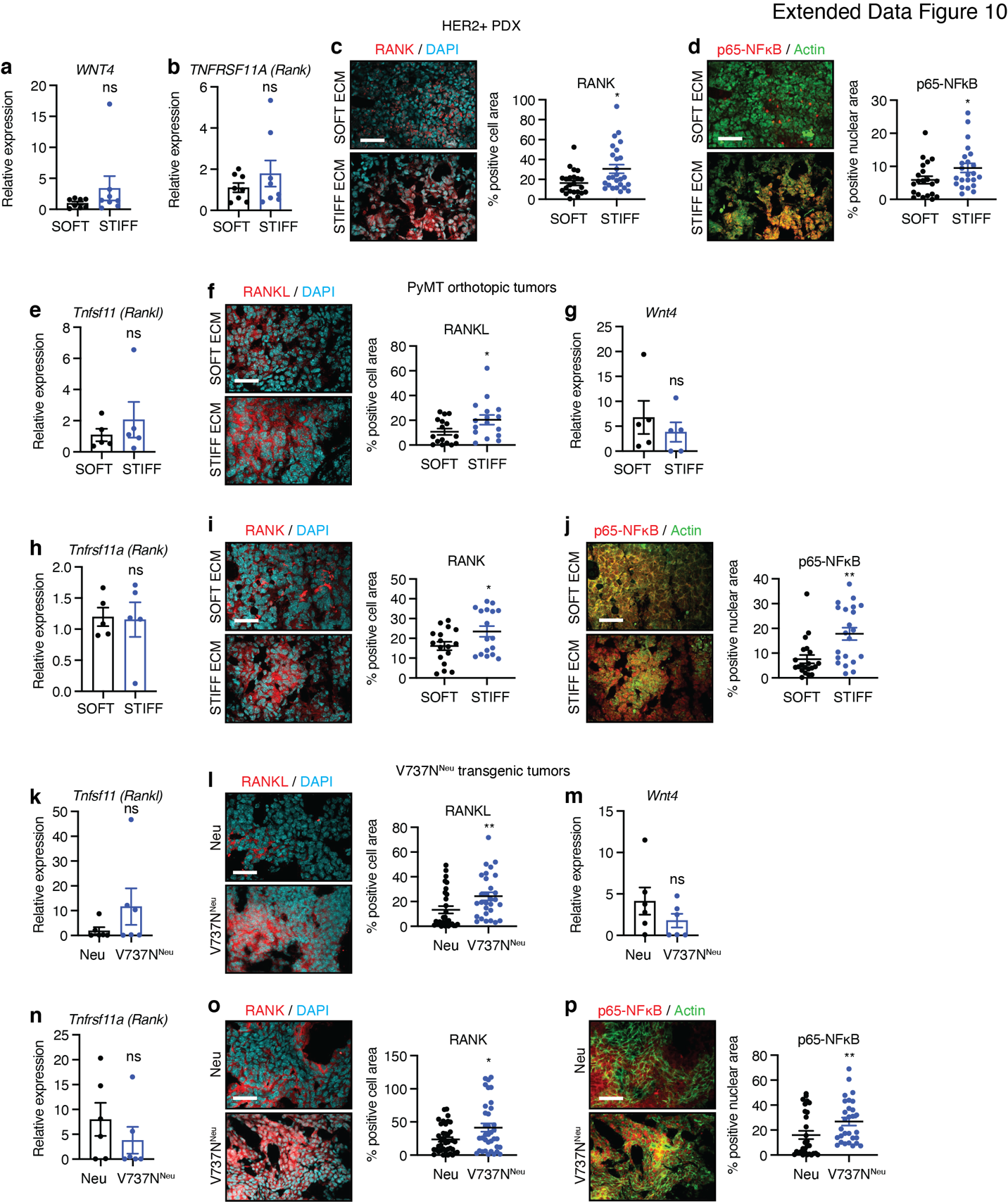
