## Supplementary figures and images for "Mechanosensitive hormone signaling promotes mammary progenitor expansion and breast cancer progression"

### Source Data

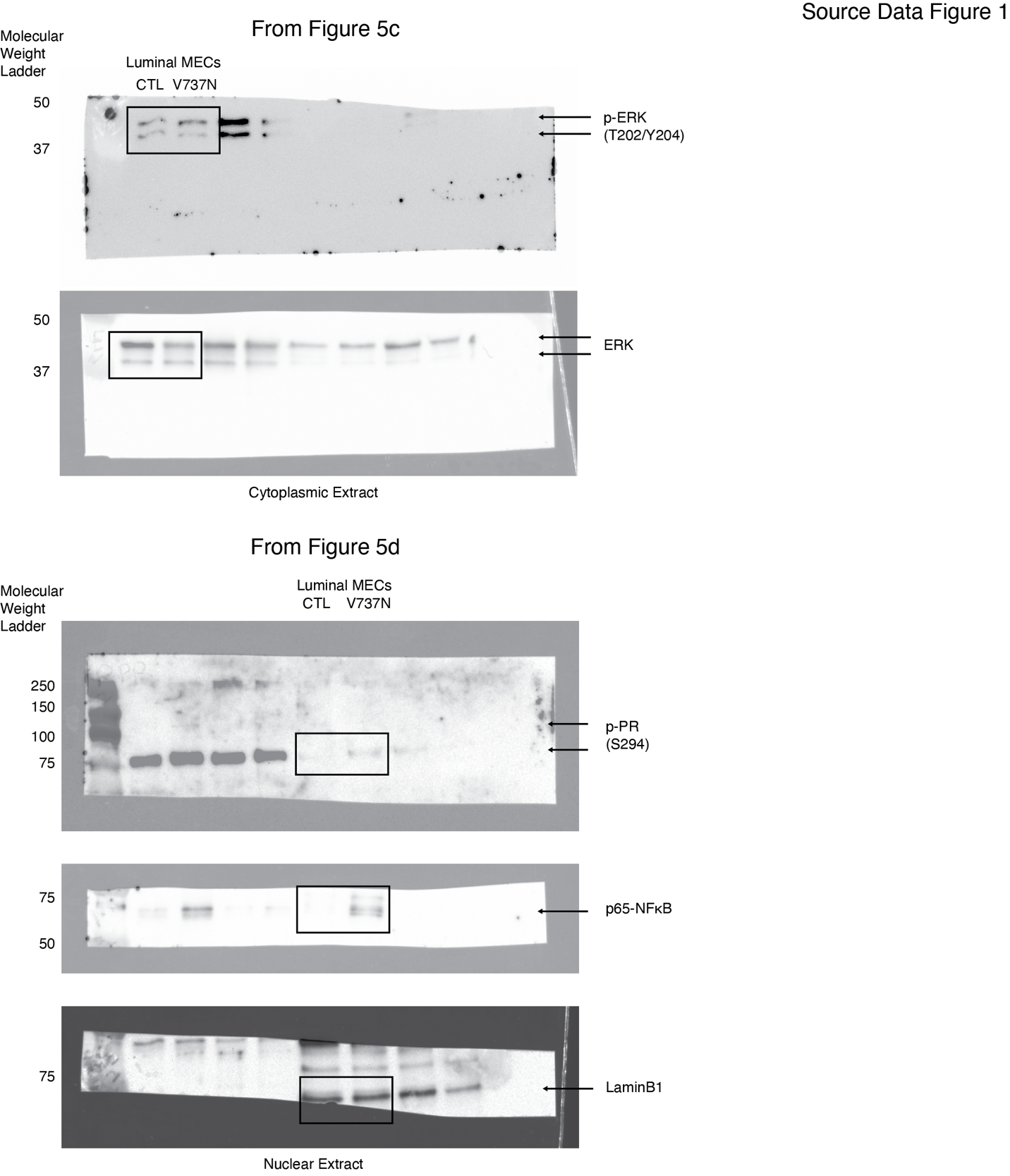

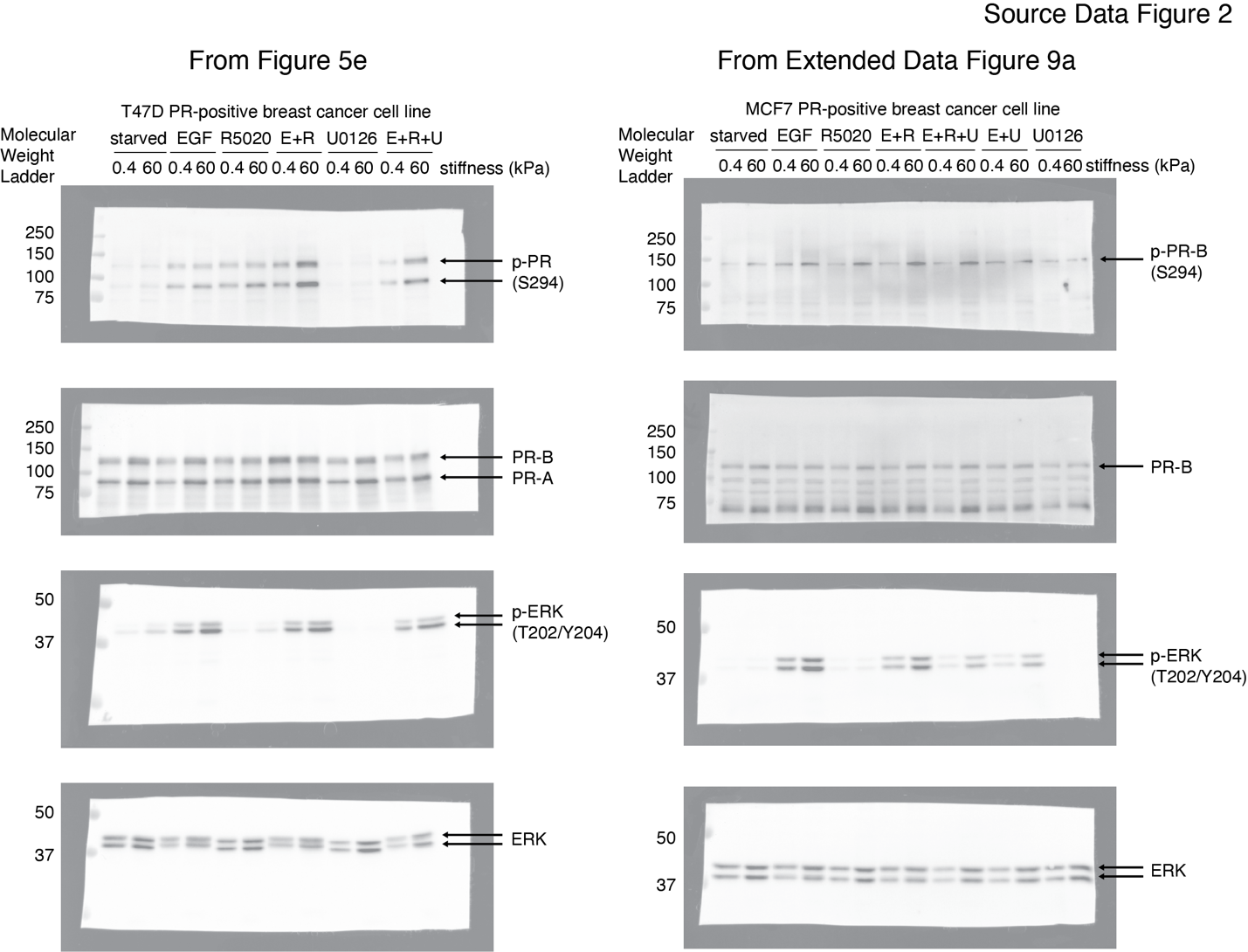

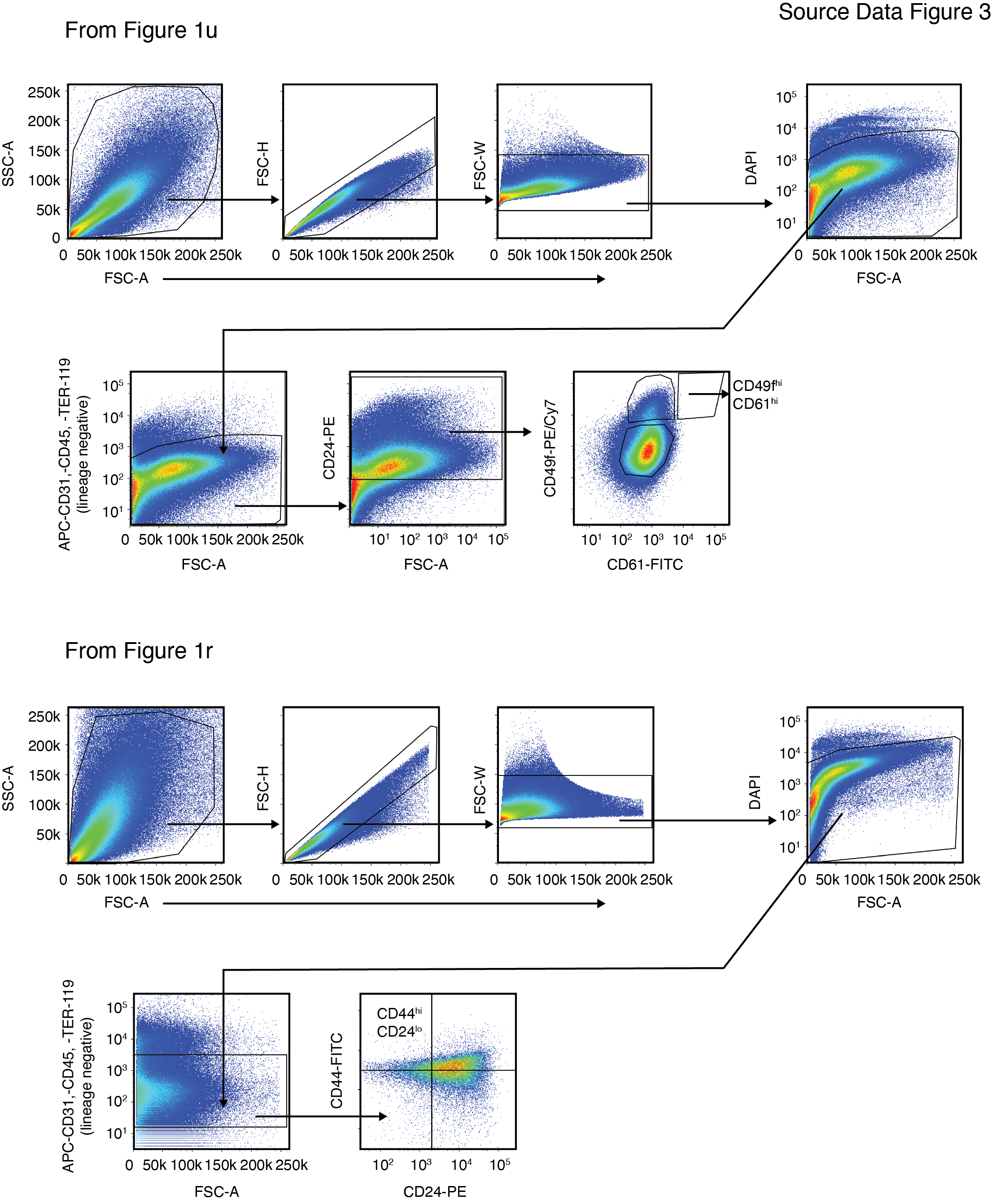

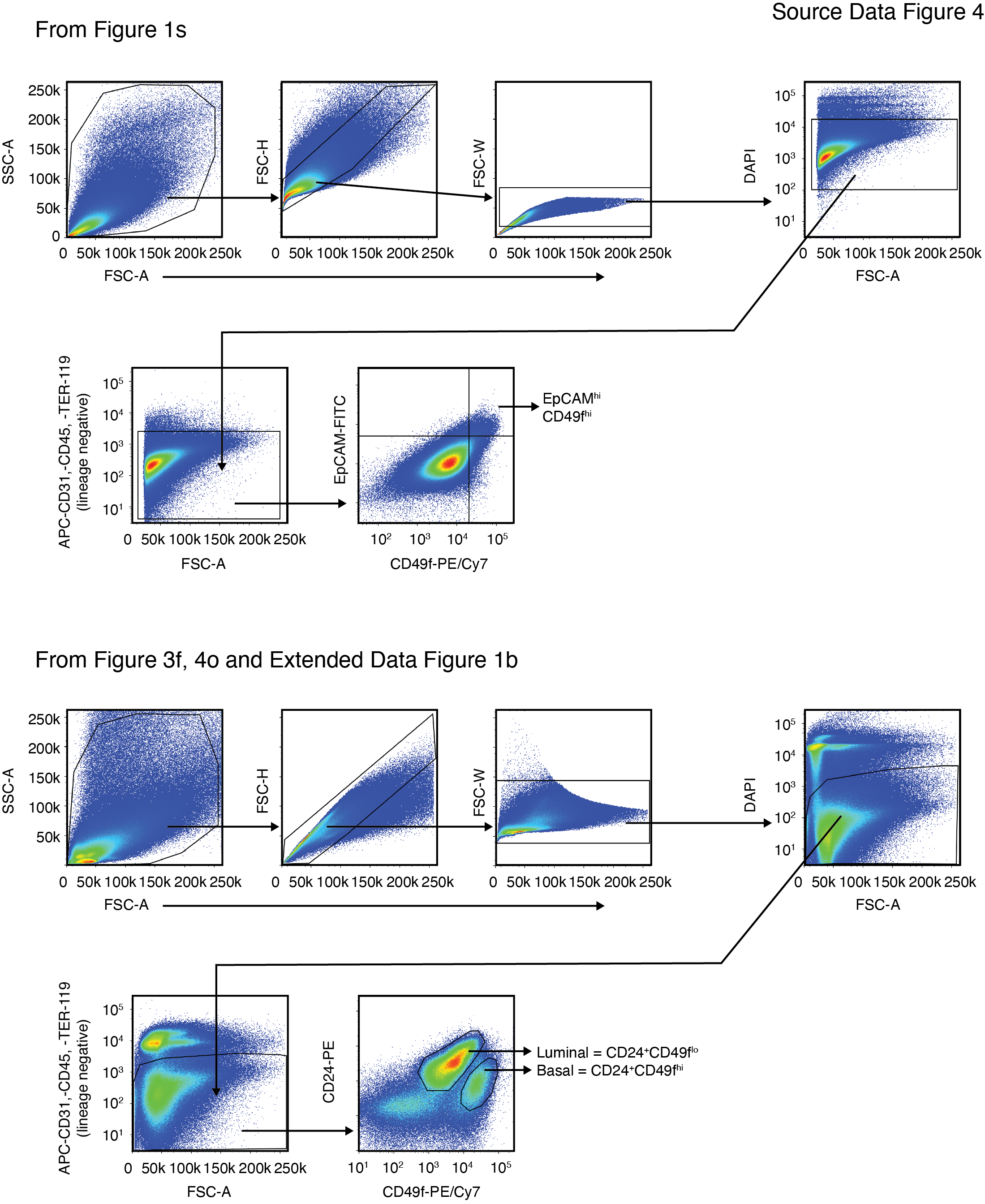

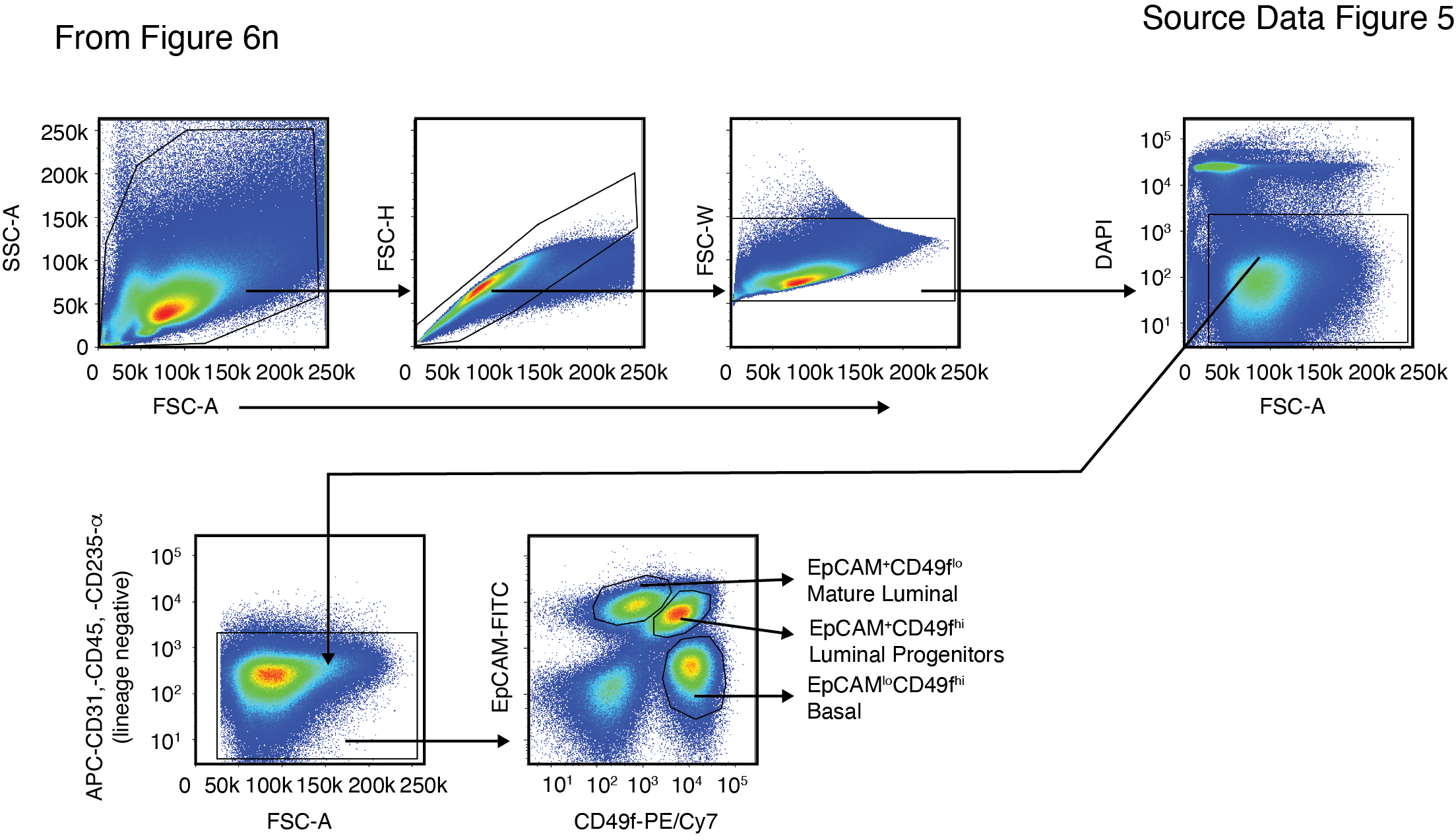
